# An examination of the evolve-and-resequence method using *Drosophila simulans*

**DOI:** 10.1101/337188

**Authors:** John K. Kelly, Kimberly A. Hughes

## Abstract

We develop a set of analytical and simulation tools for Evolve-and-Resequence (E&R) experiments and apply them to a new study of rapid evolution in *Drosophila simulans*. Likelihood based test statistics applied to pooled population sequencing data suggest parallel evolution of 138 polymorphisms (SNPs) across the genome. This number is reduced by orders of magnitude from previous studies (thousands or tens of thousands), owing to differences in both experimental design and statistical analysis. Whole genome simulations calibrated from several Drosophila genetic datasets support the contention that the observed genome-wide response could be generated by as few as 30 loci under strong directional selection, with a corresponding hitch-hiking effect. Finally, the SNPs that showed strong parallel evolution in the novel laboratory environment exhibit an (initial) allele frequency spectrum indicative of balancing in nature. These loci also exhibit elevated differentiation among natural populations of *D. simulans*.

## Introduction

In Evolve-and-Resequence (E&R) experiments, populations evolve within one or more controlled environments and are then surveyed with genomic sequencing (NUZHDIN AND TURNER 2013; LONG *et al.* 2015). The volume of data that emerges from an E&R study is remarkable, allele frequency changes at hundreds of thousands of loci within replicated populations. While researchers naturally focus on the small fraction of sites exhibiting the largest or most consistent changes, a wealth of information resides in the ‘background response’; the evolution of polymorphisms that are not direct targets of selection, which is the overwhelming majority of the genome. In this paper, we present an analytical framework for E&R studies, first to provide more detailed predictions regarding whole genome evolution, and second to robustly detect loci under parallel selection across replicate populations. We apply the method to results from a new E&R experiment on *Drosophila simulans* designed to answer two major questions: What is the genomic basis of rapid adaptation to a novel environment? What do the features of the genetic response tell us about the maintenance of polymorphisms in nature?

### The genetic basis of rapid adaptation

The traditional view of adaptive tempo is that evolution is slow relative to the ecological processes that influence contemporary populations (SLOBODKIN 1980; GILLESPIE 1991). In this paradigm, genetic change does not interact with ecological and demographic processes over the short time scale (few to several generations) encompassed by ecological processes (THOMPSON 1998; HENDRY AND KINNISON 1999; PALUMBI 2001; HAIRSTON *et al.* 2005). However, examples of rapid phenotypic evolution have been known since the mid-20^th^ century (KETTLEWELL 1958. ; FORD 1964; JOHNSTON AND SELANDER 1964) and its prevalence has become increasingly appreciated in recent years. Rapid evolution has profound practical consequences for biological control of pathogens, pests and invasive species, fisheries management, and biodiversity conservation (CONOVER AND MUNCH 2002; DARIMONT *et al.* 2009), especially in the context of accelerating climate change (WARD AND KELLY 2004). Indeed, this growing appreciation for rapid evolution has spawned new subdisciplines such as eco-evolutionary dynamics (ELLNER *et al.* 2011).

Many instances of rapid evolution of ecologically important traits have been documented in invertebrates (ELLNER *et al.* 1999; DABORN *et al.* 2002), vertebrates (REZNICK *et al.* 1997; GRANT 1999), plants (FRANKS AND WEIS 2008), yeast (LANG *et al.* 2013; LEVY *et al.* 2015), and prokaryotes (BARRICK *et al.* 2009). Biochemical (GHALAMBOR *et al.* 2015; HUANG AND AGRAWAL 2016), morphological (LOSOS *et al.* 1997; GRANT 1999), life history (ROSE 1984; HAIRSTON and WALTON 1986; REZNICK *et al.* 1997), And behavioral (TURNER AND MILLER 2012; STUART *et al.* 2014) phenotypes can evolve substantially in just a handful of generations when populations experience new selective regimes. However, less is known about the genomic changes that occur during rapid adaptation to novel environments, especially in multicellular eukaryotes (MESSER *et al.* 2016; JAIN AND STEPHAN 2017).

A key question is whether the standing genetic variation within populations is sufficient for adaptation to a novel environment, or if new mutations are required. In sexual eukaryotes, abundant standing variation is indicated by the observation that artificial selection can immediately, and often dramatically, change the mean of almost any variable trait (LEWONTIN 1974). Still, it is possible that natural selection may fail where artificial selection succeeds if the alleles that respond in artificial selection experiments are encumbered with deleterious side effects. E&R studies seem an ideal alternative to artificial selection experiments in this regard. While the researcher largely controls fitness with artificial selection, organisms “select themselves” in an E&R experiment. Pleiotropic effects on general vigor will be a major determinant of selection on alleles with favorable trait effects in an E&R experiment, but much less so in an artificial selection experiment. Admittedly, most previous E&R experiments do not evaluate the evolutionary potential of natural populations simply because they are initiated from laboratory adapted populations, or small numbers of founders. Here, we describe a replicated E&R experiment using *D. simulans* founder populations initiated with large numbers of wild-caught individuals to investigate the earliest stages of adaptive evolution.

### Genome-wide evolution in E&R studies

E&R experiments using Drosophila have addressed questions about the number and kinds of loci under selection, the relative frequency of hard versus soft selective sweeps, temporal dynamics, and the effect of selection on genome-wide patterns of diversity (BURKE *et al.* 2010; TURNER *et al.* 2011; OROZCO-TERWENGEL *et al.* 2012; REMOLINA *et al.* 2012; TURNER AND MILLER 2012; HUANG *et al.* 2014; TOBLER *et al.* 2014; KANG *et al.* 2016; BARGHI *et al.* 2017; MICHALAK *et al.* 2017; SCHOU *et al.* 2017). In a review of E&R studies, NUZHDIN AND TURNER (2013) noted a striking “excess of significance” in that thousands of polymorphisms appear to respond to selection. The number of loci that selection can act on simultaneously is an important and long-standing controversy in evolutionary genetics (HALDANE 1957; SVED *et al.* 1967; BARTON 1995). However, we generally expect that the more loci affecting fitness, the smaller the allele frequency change per locus. It is thus paradoxical that so many SNPs exhibit large change in E&R experiments. There are numerous potential reasons for excessive significance. Perhaps the simplest is that testing procedures are anti-conservative.

Hitch-hiking (MAYNARD SMITH AND HAIGH 1974) is the most likely driver of excessive significance - many significant tests are neutral SNPs in Linkage Disequilibria (LD) with selected loci (HUANG *et al.* 2014). Hitch-hiking in an E&R study requires an initial association between loci in the ancestral population(s) and also minimal subsequent recombination over the course of selection. Relevant to both, natural and laboratory adapted *D. melanogaster* populations are polymorphic for large inversions and have dramatically suppressed recombination near centromeres (CORBETT-DETIG AND HARTL 2012; KAPUN *et al.* 2014; TOBLER *et al.* 2014), potentially resulting in a large number of false positive candidate SNPs (TOBLER *et al.* 2014; FRANSSEN *et al.* 2015; BARGHI *et al.* 2017). We chose *D. simulans* for this study to evaluate genomic patterns underlying rapid evolution in a population largely free of inversion polymorphism. LD declines rapidly with the physical distance between sites in *D. simulans* (SIGNOR *et al.* 2018), but as emphasized by NUZHDIN AND TURNER (2013), sampling of haplotypes to form experimental populations can generate higher levels of LD (even at considerable physical distance) than are present in the natural population. The contribution of these sampling-generated associations to parallel evolution of replicate E&R populations can be mitigated by founding experimental replicates from distinct samples of the natural population.

### Identifying selected loci in E&R studies

A variety of analytic techniques have been developed for E&R studies, applied both at the scale of individual SNPs, e.g. BURKE *et al.* (2010), and on windows of closely linked polymorphisms, (KELLY *et al.* 2013; BEISSINGER *et al.* 2014; BEISSINGER *et al.* 2015). In some cases, tests have been catered to the specific features of the experimental design (TURNER *et al.* 2011; TURNER AND MILLER 2012; HUANG *et al.* 2014). Most frequently, the number of sequencing reads called to each alternative SNP base in each population are used as counts in a contingency table analysis; Fisher’s exact test for a single evolutionary replicate (BURKE *et al.* 2010) or the Cochran-Mantel-Haenszel test (CMH) to aggregate signal across replicates (OROZCO-TERWENGEL *et al.* 2012; HUANG *et al.* 2014; KOFLER AND SCHLÖTTERER 2014; FRANSSEN *et al.* 2015; BARGHI *et al.* 2017). As emphasized by OROZCO-TERWENGEL *et al.* (2012), contingency tables test only whether allele frequency differs between population, not whether differences imply selection. For this, researchers have employed simulations of neutral evolution to establish a threshold for the CMH statistic. Here, we implement a CMH testing pipeline, including neutral simulations, as a contrast to our likelihood/permutation method.

The genome-wide response in an E&R experiment (evolution at both selected and neutral loci) depends on the number and genomic positions of selected loci, how those loci interact to determine fitness, the nature and extent of LD, the recombination map, the reproductive biology, and the experimental design. Given these myriad factors, we do not have a clear picture of how much hitch-hiking is to be expected in a typical E&R experiment and thus a means to infer how many sites are direct targets of selection. To develop predictions, we build a simulation framework to predict the full observed response of an E&R experiment (KOFLER AND SCHLÖTTERER 2014; VLACHOS AND KOFLER 2018). The design of the experiment (how replicate populations are founded, how many individuals reproduce, and how many generations) is directly reiterated in a model that predicts change of every polymorphism in the genome. This is a parameter rich model, but prior work on *D. simulans* and its close relative *D. melanogaster* underpin essential assumptions (e.g. the recombination map, patterns of LD in nature). Observations from our specific E&R experiment establish other features such as the number and genomic positions of polymorphisms and initial allele frequencies. Finally, we extract essential information not only from the extreme outliers (putative targets of selection), but from observations on the “typical SNP” The amount and variability of change at neutral loci dispersed across the genome is an indicator of genetic draft (GILLESPIE 2001; NEHER AND SHRAIMAN 2011) and thus of selection. The simulation model provides important insights on the observed experimental results, not only in terms of the number of significant tests but also on the allele frequency spectrum at fitness determining loci.

## Methods

### I. The experiment

#### Founding populations

The ancestral populations of this experiment are from the offspring of wild-collected mated *D. simulans* females. We collected adult *D. simulans* from compost piles at Orchard Pond Organic Farm in Tallahassee, FL (Universal Transverse Mercator Grid coordinates 16N 761030 3386162) between October 28 and November 25, 2014. Adult females were isolated in vials to produce offspring. From each wild-collected female, we collected two male and two female offspring after verifying that male offspring were *D. simulans*. One male and one female offspring were flash frozen immediately to represent the founding generation (see below). These flies were kept at -80° until DNA extraction. The other male and female were used to found replicate lab populations. We initially established six replicate population cages, using one male and one female offspring of 250 wild-caught mated females per replicate and using the offspring of different wild female progenitors in each replicate. Approximately 3 weeks after founding these populations (Dec 2-3, 2014), the six cages were combined two at a time to form the A, B, and C population replicates. Equal numbers of flies were used from the pair of cages and mixed to create new cages. Thus, each of the A, B, and C populations was founded with approximately 1000 individuals descended from non-overlapping sets of 500 wild-caught, mated female parents. The rapid progression from collecting wild files to establishment of experimental populations minimized inbreeding of founders.

#### Lab rearing and maintenance

Flies were housed in plexiglass containers measuring 6028 cubic centimeters. Each cage was supplied with six 177-mL plastic bottles containing 50 mL of standard cornmeal-yeast-dextrose media. Every two weeks, we replaced three of the six bottles with bottles containing fresh media; each bottle remained in a cage for four weeks. We replaced cages with clean plexiglass containers every 28 days, in sync with a media change. No live flies were removed from cages (dead flies were removed when cages were cleaned), so populations had overlapping generations. Censuses of cages were conducted approximately every 5 weeks using digital images. Census values for each population were: A (mean=1277, range=832-1635), B (mean=849, range = 672-1147), and C (mean=1187, range=963-1620). Images were counted three times and the numbers of the three counts were averaged; these numbers might be underestimated if flies were obscured by other flies in the images.

We maintained populations A, B, and C under constant lighting and temperature conditions (12L:12D, 25°) for approximately 195 days from initial collection (population A: founded from females collected Oct 28 - Nov 1, 2014, descendants preserved May 12, 2015; B: founders collected Nov 4-Nov 11, descendants preserved May 22; C: founders collected Nov 19-Nov 25, descendants preserved Jun 5). That is, descendants of the original founders were sampled approximately 7 months after collection of the wild founders. From the last generation of each population, we collected 500 males and 500 females by aspiration. These flies were snap frozen on dry ice and kept at -80° until DNA extraction. For DNA extraction and sequencing, we pooled the 1000 preserved offspring of the founding females to form Ancestral samples A0, B0, and C0. Similarly, we pooled the 1000 flies collected at the end of the experiment to form Descendant samples (A7, B7, C7), with “7” designating months since population founding. Extractions and sequencing libraries were performed simultaneously for all six population samples.

#### Library prep and level of sequencing

We homogenized whole flies (500 males and 500 females from each Ancestral and Descendant population) and the DNA was extracted using DNAzol reagent (Thermo Fisher). We fragmented DNA using a Covaris E220 Ultrasonicator and size selected to produce insert lengths of 380-480 bp. One sequencing library was prepared for each population using the NEBNext Ultra DNA Library kit for Illumina (New England Biolabs) following manufacturers recommendations, with each population receiving a unique index. We made a distinct library from flies from Ancestral populations A0, B0, and C0 (NEB indices 13-15, respectively), and Descendant populations A7, B7, and C7 (NEB indices 16, 18, and 19 respectively). Ancestral population samples were multiplexed and sequenced in one lane, and those from the Descendant populations were multiplexed and sequenced in three additional lanes. Because one library (C7) was over-represented in the resulting data, we performed an additional sequencing run using the five remaining libraries (A0, B0, C0, A7, and B7) which were multiplexed and run on a single lane. All sequencing was conducted using an Illumina HiSeq 2500 instrument at the Translational Science Lab at Florida State University, using V3 chemistry. We sequenced 150bp on each of the paired ends. In total, DNA from 6000 flies was sequenced.

#### Sequence Analysis

We edited read pairs (fastq format) from each population sample using Scythe (https://github.com/vsbuffalo/scythe/) to remove adaptor contamination and then with Sickle (https://github.com/najoshi/sickle/) to trim low quality sequence. We used the mem function of BWA (LI AND DURBIN 2009) to map read pairs to version r2.02 of the *Drosophila simulans* reference genome, updated from the build published by HU *et al.* (2013). We used picard-tools-1.102 to eliminate PCR duplicates from the mapping files; an important step given that PCR duplicates represent pseudo-replication in bulked population samples. Prior to variant calling, we applied the RealignerTargetCreator and IndelRealigner (MCKENNA *et al.* 2010) to each population bam file. The population bams were input to Varscan v2.3.6 to call SNPs and indels. We piped the output from samtools (LI *et al.* 2009) mpileup (version 1.2) to the varscan (KOBOLDT *et al.* 2009) functions mpileup2snp (for SNPs) and mpileup2indel (for indels). We obtained the read count (number of alleles) and reference allele frequency at each variant site for each sample. We suppressed indels in downstream analyses as well as all SNPs within 5bp of an indel. We also limited attention to the major chromosomes: X, 2R, 2L, 3R, and 3L.

We scored read depths within each population prior to filtering and found that the median depth at X-linked loci was very close to ¾ the corresponding value for autosomal loci (ratio = 0.77 for Ancestral populations, 0.75 for Descendant populations). For subsequent analysis, we eliminated polymorphisms if the read depth across populations was too low for meaningful tests or atypically high across samples. For inclusion of a SNP, we required at least 60 reads per population for X-linked and at least 80 for Autosomal loci. We excluded SNPs if the total read depth within Ancestral and Descendant (considered separately) was greater than the 95^th^ percentile of the corresponding depth distribution. Here, we conducted separate filtering on autosomal sites and X-linked sites as the latter have lower coverage. After filtering, 291272 SNPs remained (58647, 49940, 69010, 71289, and 42386 on 2L, 2R, 3L, 3R, and X, respectively). The read depth and allele frequency at each SNP in each population is reported in Supplemental Table S1.

#### Comparison to other datasets

We attempted to ascertain the loci investigated in this experiment within two other recent sequencing studies of *D. simulans*. MACHADO *et al.* (2016) and SIGNOR *et al.* (2018) also mapped reads to the w2 reference published by HU *et al.* (2013). However, each of the three studies used different versions of this genome build with different coordinates for homologous sites: version r2.01a for MACHADO *et al.* (2016), version r2.01b for SIGNOR *et al.* (2018), r2.02 for this paper. To compare loci, we extracted the 100bp sequence containing each of our 291,272 SNPs in the r2.02 reference genome and mapped these sequences to the two previous genome builds (using BWA and Samtools as described above). In most but not all cases, the r2.02 sequence mapped to a single unique location in the r2.01a and r2.01b genomes. Comparing homologous sites, we find that polymorphism observed in one study are often not reported in one or both of the others. This may be biological (if one of the alleles is fixed in the other populations) or bioinformatic (the polymorphism is present but not ascertained in the other populations), but the latter is clearly a major factor. SIGNOR *et al.* (2018) report many more polymorphisms than MACHADO *et al.* (2016) or this study, despite sequencing a much smaller collection of flies (170 inbred lines), probably because variants are more confidently called in a collection of individual inbred lines, each sequenced to high depth, than in pooled population samples. Despite these issues, it is reassuring that when we do observe that the same polymorphism across samples, the same bases are typically identified as alternatives in each population. This is demonstrated for a relevant set of loci in Supplemental Table S2.

MACHADO *et al.* (2016) surveyed 8 natural populations of *D. simulans* over a geographic transect (Florida to Maine). To test whether ‘selected loci’ from the present study were more or less geographically divergent than neutral loci, we determined the mean F_st_ (WRIGHT 1943) within windows around our significant loci (see RESULTS) and compared this value to a null distribution established by randomly sampling windows from the genome as a whole (all ascertained loci). In this context, an ascertained locus is the sequence flanking each of our 291,272 SNPs (r2.02 build) that can be uniquely and perfectly matched to a sequence in the main chromosomes of the r2.01a build. Given a set of SNPs, we calculated F_st_ at each site based on the counts of reads for each allele in each population. We excluded sites that were not scored in all 8 populations or had fewer than 125 reads in total. When windows overlapped, a SNP was only used once. Scoring individual reads as binary variables (0 if reference, 1 if alternate), the one-way ANOVA provides an unbiased estimate for the within and among group variance. F_st_ is the ratio of among group to total variance. We then averaged F_st_ across all sites in that specific dataset. We conducted these calculations for a range of window sizes.

### II. Analysis of evolutionary change at SNPs

#### Null divergence

The raw data for each polymorphism is twelve numbers, the counts of reads for each alternative SNP base in each of the six populations (A0, B0, C0, A7, B7, C7). The statistical treatment of data is based on transformed allele frequencies (FISHER AND FORD 1947; WALSH AND LYNCH 2018). If *p*_*A*_ and *p*_*D*_ are allele frequencies in Ancestral and Descendant populations, respectively, then 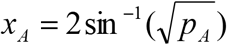 and 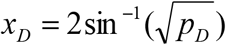, with x measured in radians (the A/D subscript denotes Ancestral and Descendant, respectively). This angular transformation is useful because, to a first approximation, the variance in change owing to genetic drift and sampling is independent of the true allele frequency (FISHER AND FORD 1947). As a consequence, a common test can be applied across polymorphisms despite differing initial allele frequencies (KELLY *et al.* 2013). At a neutral autosomal SNP, divergence (*x*_*D*_ - *x*_*A*_) is normally distributed with mean 0 and variance 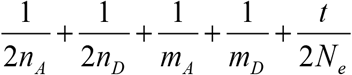, where m = read epth at the locus after sequencing, n = number of diploid individuals sampled for sequencing, N_e_ is the effective population size, and t is the number of generations. While five of these quantities are known, N_e_ is not. However, we can use the observed variance in divergence to estimate N_e_. To estimate N_e_, we focus on autosomal loci given that N_e_ differs between autosomes and the X (CHARLESWORTH 2009), and 85% of our SNPs are on the autosomes. We estimate the “null variance”, ***ν*** on a population and chromosome specific basis using:

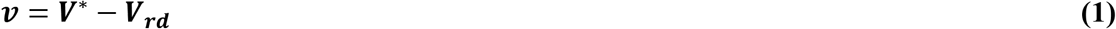

where *V*\* is a robust estimator for the variance in divergence between Ancestral and Descendant populations and *V*_*rd*_ is the read depth variance. *V*\* is calculated from the interquartile range of the distribution of changes in allele frequency (KELLY *et al.* 2013). *V*_*rd*_ is estimated from the average of 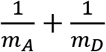 across all SNPs. ***ν*** aggregates the dispersive processes that are shared by all neutral SNPs; stochastic changes in allele frequency over the course of the experiment as well as the sampling of flies into bulks and any differential representation of the DNAs from each fly within the DNA pool. To estimate ***ν*** for each chromosome, we focus on SNPs with 0.05 < *P*_*A*_ < 0.95 to avoid boundary effects. In the neutral case,

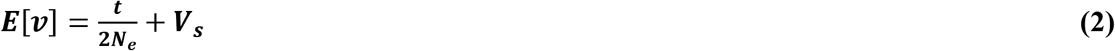

where *V*_*s*_ is the total variance associated with bulk formation. With selection, ***ν*** may be determined as much (or more) by genetic draft than by genetic drift.

#### Tests for selection

Divergence between Ancestral and Descendant populations is statistically independent between replicates because each population was founded from distinct flies and there was no gene flow between replicates. Under the null hypothesis, *E*[*x*_*A*_ - *x*_*D*_] = 0 and the variance is given by *v*, *m*_*A*_, and *m*_*D*_. The likelihood for the null model at each SNP (LL_0_) is the product of three normal densities. We contrast the normal density likelihood of the data under this model to an alternative allowing parallel evolution across replicates: *E*[*x*_*D*_ – *x*_*A*_] = δ, where δ is the (shared) change in allele frequency due to selection across replicate populations. For the alternative model, we also require the MLE of 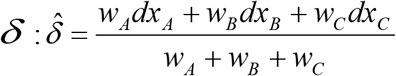 where the *dx* terms are the observed (*x*_*A*_ – *x*_*D*_) in each replicate and the are replicate specific weights: 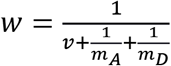(see MONNAHAN AND KELLY (2017) for derivation). The Likelihood Ratio Test (LRT-1) for parallel evolution is 2(LL_1_-LL_0_). This is not a strict likelihood in the sense that v are treated as constants and not free parameters, but the error associated with this procedure is minimal given that v are estimated from the aggregate of thousands of variant sites (MONNAHAN AND KELLY 2017). Also, we use permutation (and not the parametric chi-square distribution) to assess genome-wide significance LRT-1. TO create a permutation replicate, we randomly scrambled observed standardized divergences across SNPs within each replicate population and then calculate LRT-1 at each SNP. The distribution of divergences (across SNPs within each population) is preserved by this procedure and so the test is based entirely on consistency of response of the same SNP across replicates. We extracted the largest LRT-1 value from each permuted dataset and repeated the procedure 1000 times.

A second likelihood ratio, LRT-2, tests for heterogeneous divergence among populations. The most general model allows the expected change, *E*[*x*_*D*_ – *x*_*A*_], to be population specific. There are three free parameters in this model, a distinct *δ* for replicates A, B, and C. The MLE for the *δ* of each population is simply the observed *x*_*D*_ — *x*_*A*_. LRT-2 is 2(LL_2_-LL_1_) which we compare to a chi-square distribution with 2 degrees of freedom. Permutation cannot be used for LRT-2 because the null hypothesis (selection is consistent across replicates) is not reiterated by randomizing SNP locations (Table 1).

**Table 1.** A summary of the tests introduced in this study.

| Test statistic | Signal to be measured | Null distribution |
| --- | --- | --- |
| LRT-1 | Parallel selection at individual SNPs | Permutation of SNPs or $\chi^2$ with 1 df |
| LRT-2 | Heterogeneous selection at individual SNPs | $\chi^2$ with 2 df |
| Extreme windows | Loci with extreme response across replicates | Permutation of windows |
| Window rank correlation | Consistency of response across replicates | Permutation of windows |

We apply two additional tests at the scale of genomic windows. LRT-1 significance requires the same SNP to show a strong parallel response, which may not occur at a selected site because allele frequency estimates are encumbered with substantial error. Closely linked sites that all respond to selection owing to hitch-hiking will exhibit differing signals across replicate populations owing to differing estimation error. To capture the selection signal at such loci, we first identified, within each replicate, the site in each window with the maximum absolute value of (standardized) divergence between ancestor and descendant. We constructed non-overlapping windows, each with 25 adjacent SNPs, and assigned that maximum value to the window. We then used the sum of window specific scores across replicates as a test statistic - loci exhibiting a strong response across replicates will have large sums (Extreme windows test in Table 1). We established a significance threshold by permuting entire windows, preserving the linkage relationships among neighboring sites. The selection regimes that are distinctly measured by LRT-1 (parallel selection) and LRT-2 (heterogeneous selection) could both contribute positively to Extreme windows test.

Finally, we also calculated the Spearman Rank correlation (R) of window scores in pairwise contrasts between populations (A vs. B, A vs. C, and B vs. C). The average R measures the general tendency for magnitude of change per locus to vary consistently across independent replicate populations. Unlike the first three tests, it is not based on outliers. The permutation procedure for Extreme windows is suitable here, except with the R values recalculated with each permuted dataset and entire windows permuted. We first applied the Window rank correlation test to the entire dataset (about 11,500 windows) and then to a more restricted collection of windows (about 8,500). The latter excluded regions near centromeres and telomeres (low recombination) and also excluded windows that were diagnosed as significant by the Extreme windows test.

### III. The Simulator

The program tracks each chromosome of each individual in the population as a series of binary values (the allele present at each locus). The position of each of the 291,272 SNPs in the simulation matches positions observed in the experiment. To optimize use of computer memory, the program compresses genotypic information using the method of SUKUMARAN AND HOLDER (2011). The population is defined as N_m_ male and N_f_ female diploid adults formed each generation (only one X haplotype in males). Each subsequent generation is formed by randomly selecting parents and synthesizing gametes from those parents to create a new set of N_m_ males and N_f_ females. In simulations with selection, individuals are chosen with probabilities proportional to their fitness (which is a function of the genotype at the subset of ‘selected’ loci). We first consider neutral evolution and then two different models of selection. In the truncation selection model, individuals have fitness 1 if their genetic score (a sum of effects across loci) exceeds the threshold and 0 otherwise. In the multiplicative model, each site affecting fitness has a selection coefficient (s) of 1, 1+s, or 1+2s. Individual fitness is a product across loci. For both selection models, we assume that hemizygous male genotypes have the “homozygous” effect, e.g. 1 or 1+2s. A simulation replicate starts with creation of the founding population (described below) followed by sampling of three distinct experimental populations, each propagated for 15 generations (selection occurring in 14 of those generations if indicated). In each population, we simulate data collection to mimic the actual experiment with read depths at polymorphic sites equivalent to the observed values.

Two challenges for a system-specific forward simulation are (1) how to synthesize the “founding haplotypes”, i.e. the genome sequences of generation 0 individuals, and (2) how to impose recombination events in a realistic manner along chromosomes. Regarding (1), the sequence data from our pooled generation 0 samples (A0, B0, and C0) provide strong information about allele frequencies, but not about haplotype structure in the ancestral natural population. Strong information about haplotype structure is provided by the inbred lines sequencing of SIGNOR *et al.* (2018), but this is for a different population of *D. simulans.* LD between alleles is often idiosyncratic to population, and for this reason, we do not extract specific LD values from the SIGNOR *et al.* (2018) lines. Instead, we use these sequences (downloaded from https://zenodo.org/record/154261#.W2hyqhhKhrk) to estimate the overall strength of association between loci, or more precisely, the probability distribution of linkage disequilibrium conditional on the frequencies of alleles at each locus and the distance between sites (Supplemental Table S3; python scripts included in Supplemental file 1). We established distinct probability tables for autosomes and the X chromosome (see Table 1 of SIGNOR *et al.* (2018)) and also noted elevated LD in low recombination regions of the genome (near centromeres and telomeres; Supplemental Table S4; Supplemental Figure S1). We separated these regions and calculated LD probability tables based on the “normal recombination” regions of the X and autosomes (Table S3). However, to simulate LD within reduced recombination areas, we adjusted the probabilities to produce average LD values similar to those observed in low recombination regions in the real data. We found that increasing the probability of the lowest value for D’ (LEWONTIN 1964) by 10% and reducing the probability of all intermediate values by 20% yielded a good match between simulated and observed values for average r^2^ versus genomic distance (Table S2).

The frequencies of alternative SNP bases and the locations of polymorphisms are specific to our ancestral population (estimated from the A0, B0, C0 data). The founding population of a simulation replicate is synthesized by stochastically sampling LD values conditional on allele frequencies and locations. The three experimental replicates within a simulation replicate are sampled from the same founding population, but different simulation replicates will have distinct founding populations owing to the stochastic determination of LD. Given a founding population, we sample the haplotypes of each animal in each replicate population as a vector of 0s and 1s (reference or alternative base at each SNP). We use a ‘target locus’ approach for generating this vector, which is the length of the number of SNPs on each chromosome arm. 238 of the 291,272 SNPs are denoted as target loci, located at approximately 500kb intervals across the genome. In simulations with selection, the 30 fitness determining loci are a subset of the target loci. To simulate a haplotype, we first randomly sample alleles at target loci given site specific allele frequencies. We assume no LD between target loci so these samples are independent. Given the allele at a target locus, we fill in SNPs sequentially by moving out from each target locus. The remaining 291,034 SNPs are “partners” to a specific target locus (the closest one) which is always within 250kb. The probability of obtaining a particular allele (0 or 1) at a partner site is determined by the allele frequency at that site, the identity of the allele at its target locus, and the LD between these sites (determined previously when the founding population was simulated).

We use the target locus approach to generate realistic patterns of associations between targets and partners given that the full vector for a chromosome is not accurately described as a Markov Chain. Table S1 indicates much greater LD is observed between distant SNPs than is obtained if one simulates a chromosome by progressing sequentially and conditioning only on the last SNP. Our simulations with selection specify the fitness determining loci as a subset of the targets. We thus reiterate the linkage between selected and neutral loci that is essential to hitch-hiking. However, the target locus approach may fail to accurately describe (and likely underestimate) associations between closely linked loci that are not target-partner.

The simulation of gamete formation depends on the overall rate of cross-over per chromosome and the locational distribution of these events. We assume no recombination in males. TRUE *et al.* (1996) provide reasonable estimates for the map length (recombination probabilities) per chromosome arm in *D. simulans* females: 2L: 0.570, 2R: 0.655, 3L: 0.555, 3R: 0.830, and X: 0.590. The location of crossovers, when they occur, is probabilistic. Fine scale recombination data is available from *D. melanogaster* (COMERON *et al.* 2012), but not yet for *D. simulans.* For the present study, we simulate recombination given the overall rates from TRUE *et al.* (1996), but use the location distribution from *D. melanogaster* (COMERON *et al.* 2012), which exhibits reduced recombination rates near telomeres and centromeres. Recombination probabilities as a function of genomic location (500 kb intervals) are reported in Supplemental Table S5. This component of the simulation will be improved when fine scale recombination data from *D. simulans* becomes available.

For each parameter set (specified values for N_m_ and N_f_, the number and location of selected sites, selection coefficients at each site, etc.), we simulate the entire experiment 1000 times to determine the range of outcomes. We then subject the simulated data from each replicate to the same analysis pipeline as applied to the real data. We first consider neutral evolution to confirm the validity of eqs (1-2) for estimating N_e_, and also to provide the null distribution for the CMH tests that are subsequently applied to both real and simulated data. We next consider a range of cases with both truncation and multiplicative selection. We evaluate these outputs in relation to multiple aspects of the observed results including the magnitude of allele frequency change at selected loci, the mean and variability in the null variance (ν) across chromosomes and replicate populations, and the distribution of test results obtained by different testing procedures. The simulation programs were written in C and are included in Supplemental File 1.

#### Data Availability Statement

Supplemental files available at FigShare. Table S1 summarizes responses at the full set of 291272 SNPs. Table S2 is the SNPs significant for LRT-1 where a polymorphisms was also observed in the other *D. simulans* experiments. Table S3 is the two locus haplotype frequencies conditional on allele frequencies and the distance between loci. Table S4 is a summary of LD estimates with varying distance. Table S5 is the crossover location distribution used for simulation. Table S6 is the 138 polymorphisms with an LRT-1 value (test for parallel selection) that exceed the permutation threshold of 47.0. Table S7 is the maximum LRT-1 value (genome-wide) for each of 1000 permutations of the data. Table S8 is all SNPs that were significant by either LRT-1 or CMH. Figure S1 reports LD measured as r^2^ as a function of chromosomal location from SIGNOR et al. (2018). Figure S2 is the average Spearman correlation obtained by permutation of windows and from simulations with selection after excluding windows in low-recombination regions and those containing LRT-1 significant SNPs. Figure S3 is the distribution of significant LRT-1 tests per experiment for 1000 simulations of the base parameter set. Supplemental.File.1 is a bundle of code. Sequence data will be submitted to the NCBI trace archive upon manuscript acceptance.

## Results

138 polymorphisms are genome-wide significant for LRT-1 indicating parallel adaptation (Figures 1-2, Supplemental Table S6). The LRT-1 value for these SNPs is larger than the most extreme single test in 95% of the permuted datasets (47.0 is the permutation threshold; Supplemental Table S7). Importantly, these sites do not all represent distinct “evolutionary events” given that significant variants are often closely linked. Figure 1 illustrates the largest LRT-1 value per 100kb window along each chromosome. There is clearly an aggregation of strong signal in regions of low recombination. The clumping of LRT in these regions nicely parallels the pattern of high LD among distant sites (Supplemental Figure S1). If we “bin” significant tests that are closely linked (within 1 mb), then 30 distinct “loci” are evident as putative targets of selection across the genome (gold triangles in Figure 1). There was only one genome wide significant test for LRT-2 (X chromosome position 14386563, p = 2.81×10^−9^) and the Extreme window test yielded only 21 significant loci, all of which contain SNPs that were significant for LRT-1 (Figure 1; Table S9). The Window rank correlation is significantly positive. With all windows included (Figure 3), the average R across pairwise contrasts of populations is 0.11; much greater than any value obtained by permutation. If we exclude excessively divergent windows or those in recombination suppressed regions, the average R is reduced to 0.061 but remains highly significant (p < 0.001; Supplemental Figure S2).

**Figure 1.**
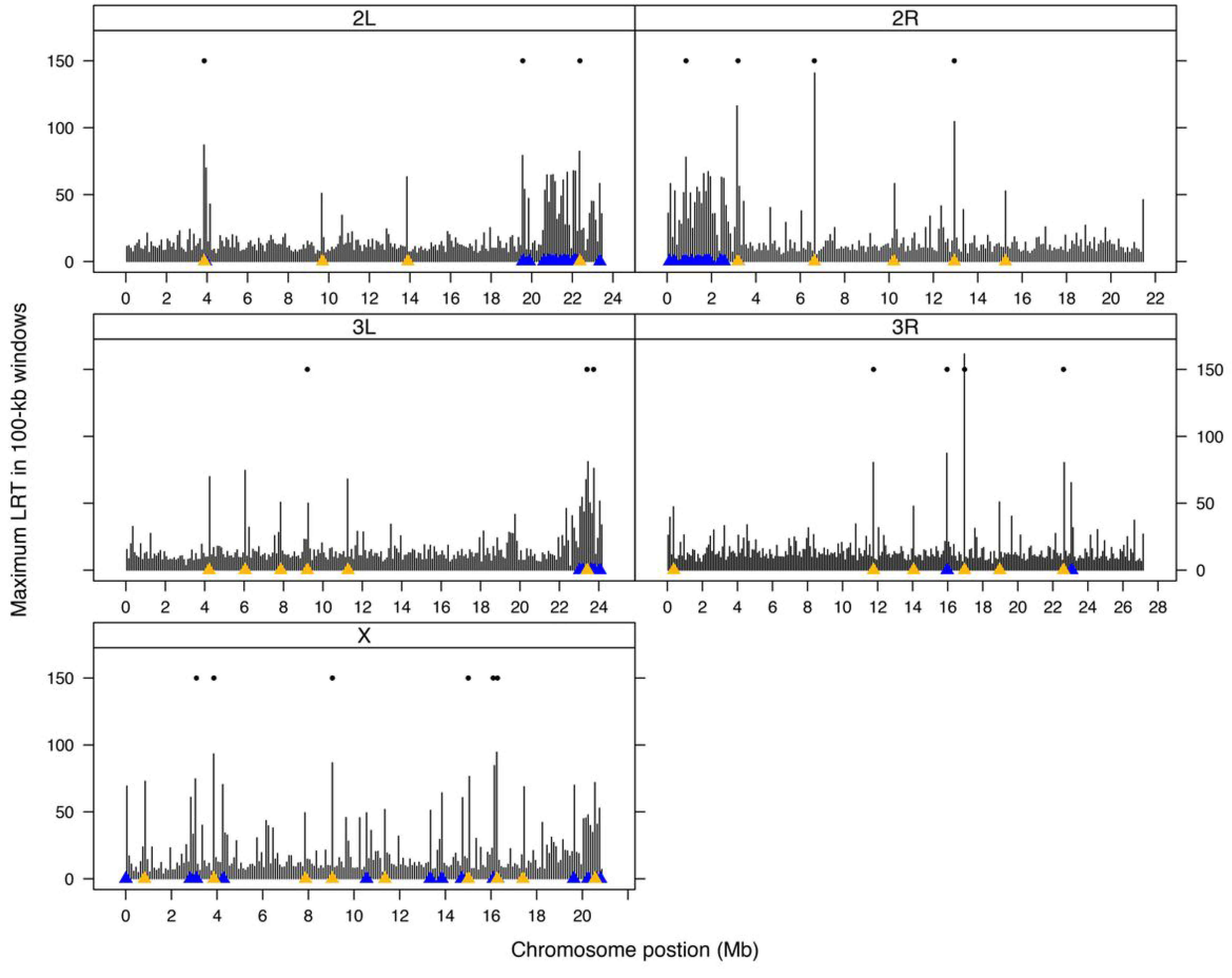
The maximum LRT-1 value (parallel selection) per 100kb window along each chromosome is indicated by vertical bars. Triangles (x-axis) indicate genome-wide significant LRT-1 tests with gold denoting the most significant test per ‘locus.’ The floating dots indicate windows significant for parallel change.

**Figure 2.**
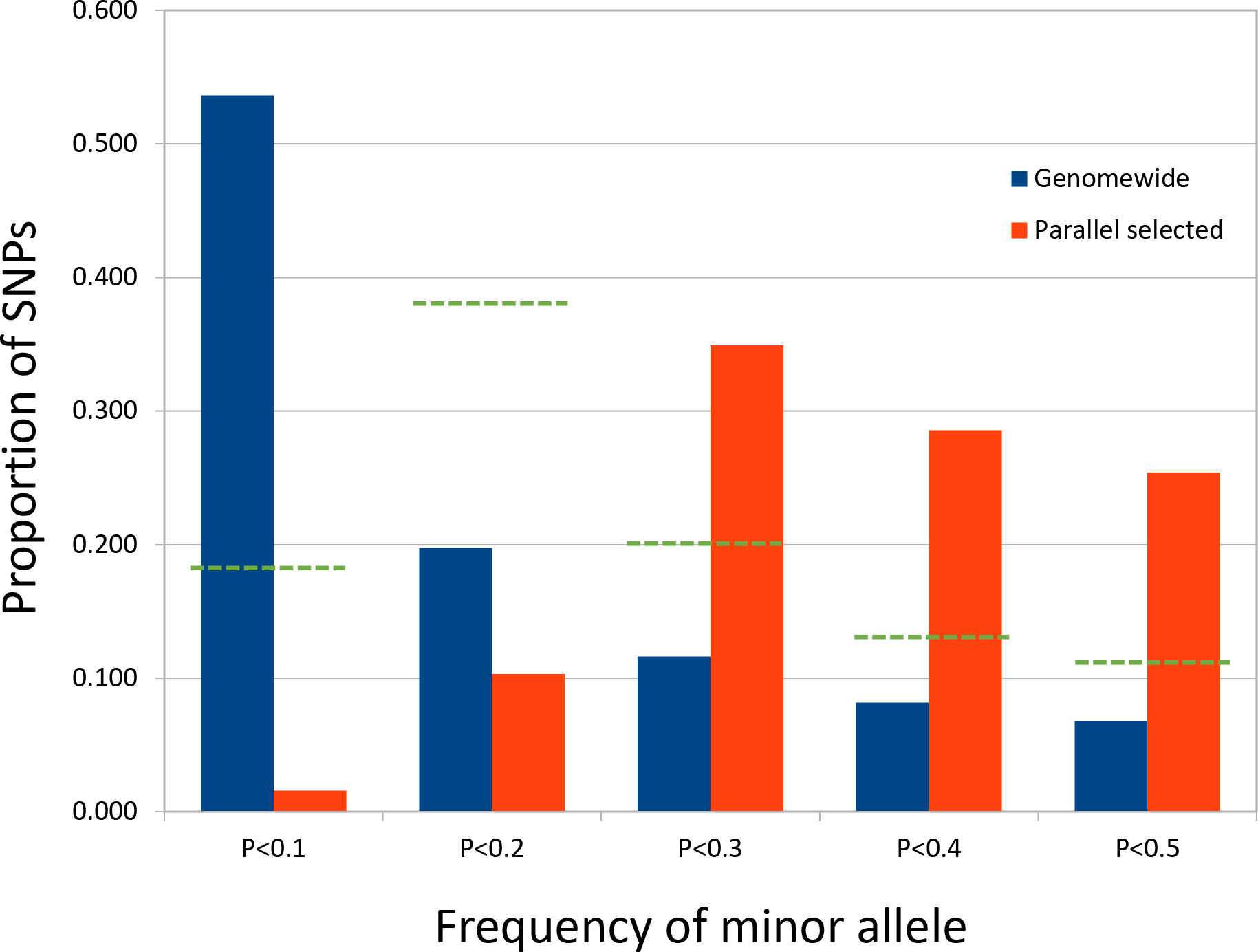
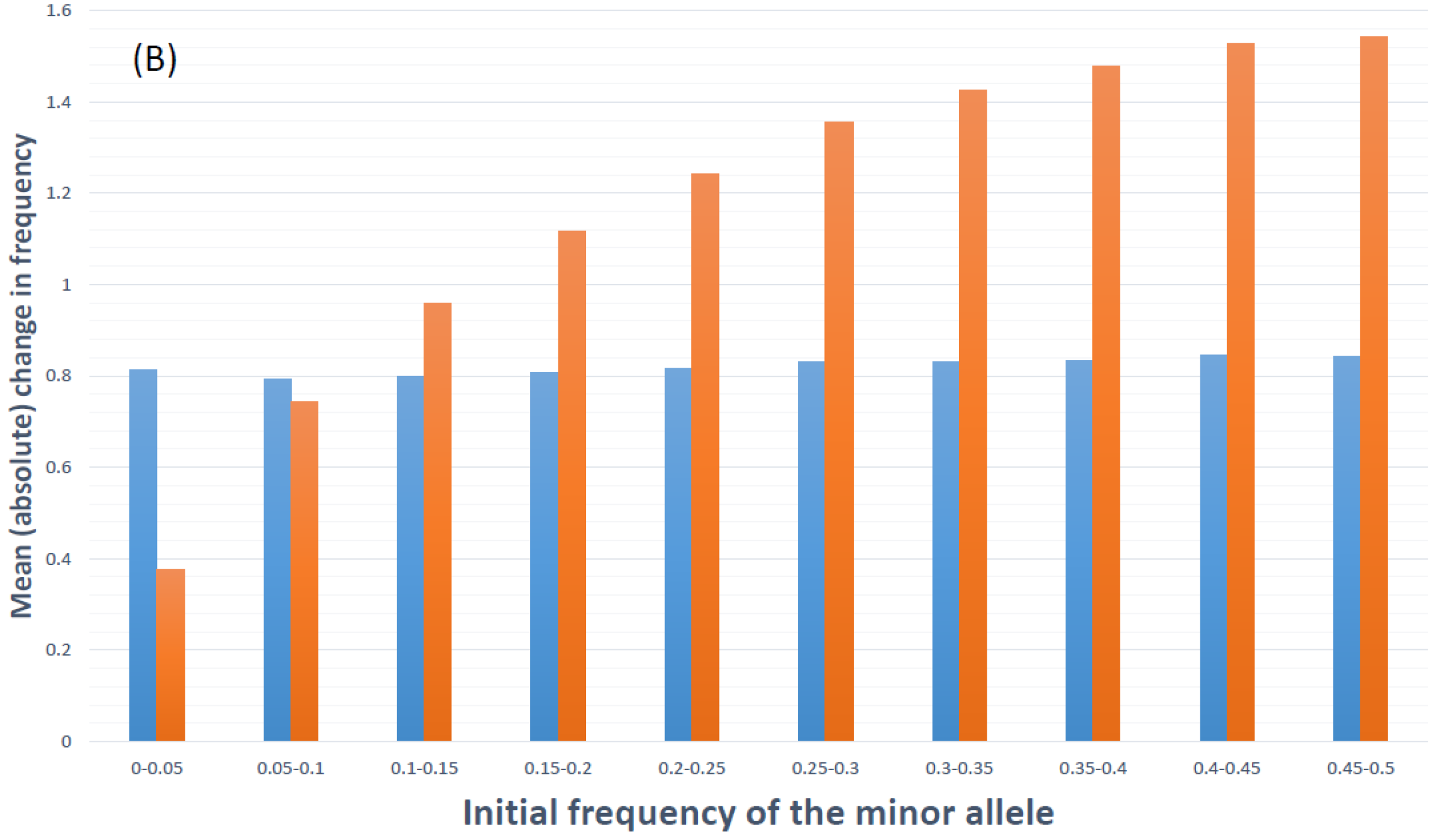
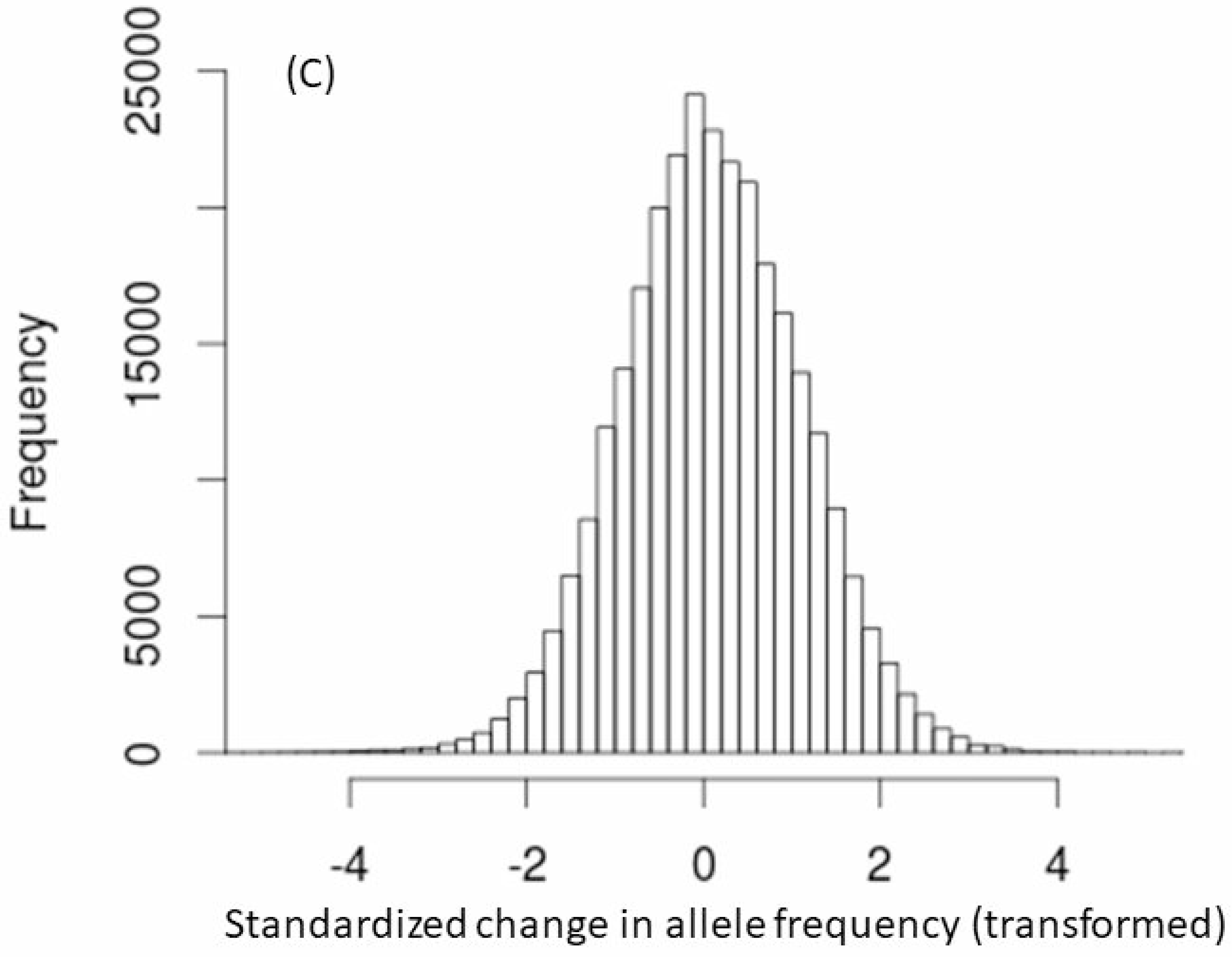
(A) The allele frequency spectrum is given for all SNPs in the ancestral populations (blue bars) and for those that tested significant for parallel selection (orange bars). The green broken lines denote proportions observed in a simulation of parallel selection on 30 loci with initial allele frequencies sampled from genome wide distribution (see text). (B) The average absolute change in z (blue bars) and of untransformed allele frequency (orange bars) as a function of initial allele frequency. The latter is multiplied by 20 to be on same scale as the former. (C) The distribution of changes in z between Ancestral and Descendant populations of replicate C (all SNPs). For (B) and (C), z is the difference in transformed allele frequency (*x*_*A*_ - *x*_*D*_) divided by the standard deviation.

**Figure 3.**
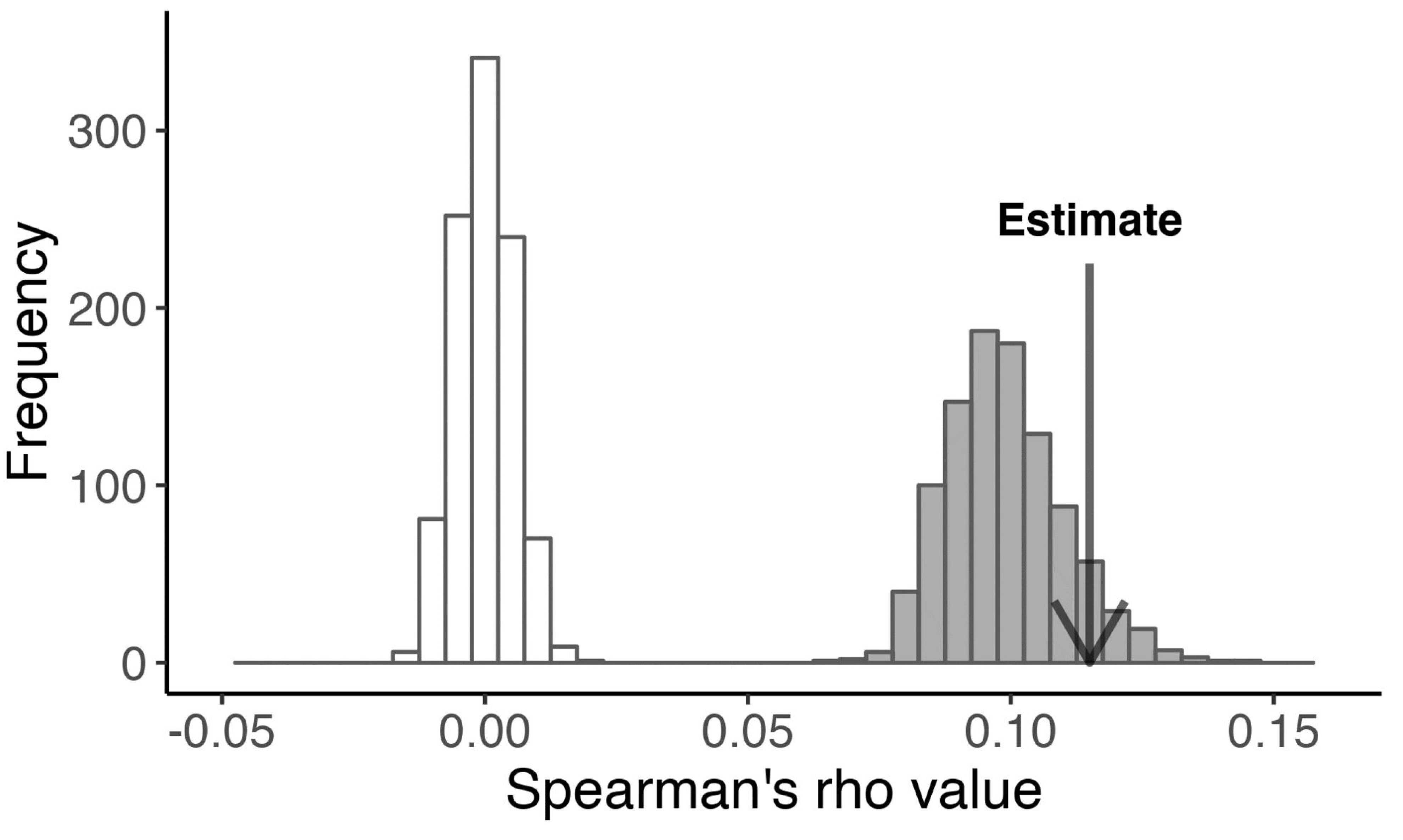
The average Spearman correlation is indicated relative to the distribution obtained by permutation of windows (white bars) and from simulations with selection on 30 loci (filled bars = base parameter set). These correlations were calculated with all windows included.

The striking feature of selected SNPs identified by LRT-1 is that nearly all have intermediate frequencies in the natural population (Figure 2A). The overall allele frequency spectrum (AFS) in the ancestral population (blue bars) is typical of natural population samples of *D. simulans* (SIGNOR *et al.* 2018). Less than half of SNPs have a minor allele frequency (MAF) greater than 0.1. However, among SNPs testing positive for parallel selection (orange bars), 98% have MAF > 0.1 in the ancestral population and 88% have MAF > 0.2. This pattern is unchanged if we thin the data by taking only the most significant SNP within each of the 30 loci remaining after binning (see above). Among these, the MAF is less than 0.1 for only one SNP and greater than 0.2 for 26 of 30.

Large change at intermediate frequency polymorphisms is not a general feature of the data set as a whole. Across all loci, divergence of transformed allele frequency is essentially homoscedastic, although this is not true of untransformed frequencies (Figure 2B). The overall distribution of changes in transformed frequencies is also highly normal (Figure 2C) justifying use of the angular transform (FISHER AND FORD 1947). The null variance in divergence (***ν***) varies substantially among chromosomes and replicate populations (Table 2) with an autosomal average of ***ν*** = 0.0185. We can estimate N_e_ using eq (1): 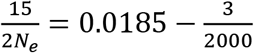, which yields N_e_ = 440.9. Here, 15 is the estimated total number of generations (14 of selection plus the production of one progeny generation, assuming a 13-day generation time) and 3/2000 is *V*_*s*_, the estimated variance owing to three sampling events, each involving 1000 diploid flies (from ancestrally sampled females to Ancestral DNA pool, from ancestrally sampled females to generation 0 of the experimental population, and from the final generation of the Descendant population into the Descendant DNA pool). We are ignoring any differential representation of individual fly DNA within the pools and also the unknown relatedness of flies between Ancestral pool and Ancestral population 0 (which depends on number of sires per wild caught pregnant female). Accommodation of these factors might cause slight changes in the estimated N_e_ but would not affect the LRT tests for selection. Each of the tests in Table 1 depend only on ***ν*** and not on the underlying components (drift versus bulk sampling variance). The numerical estimate for N_e_ is used here only to establish a null distribution for the CMH test when we contrast it to LRT-1 (see simulation results below).

**Table 2.**
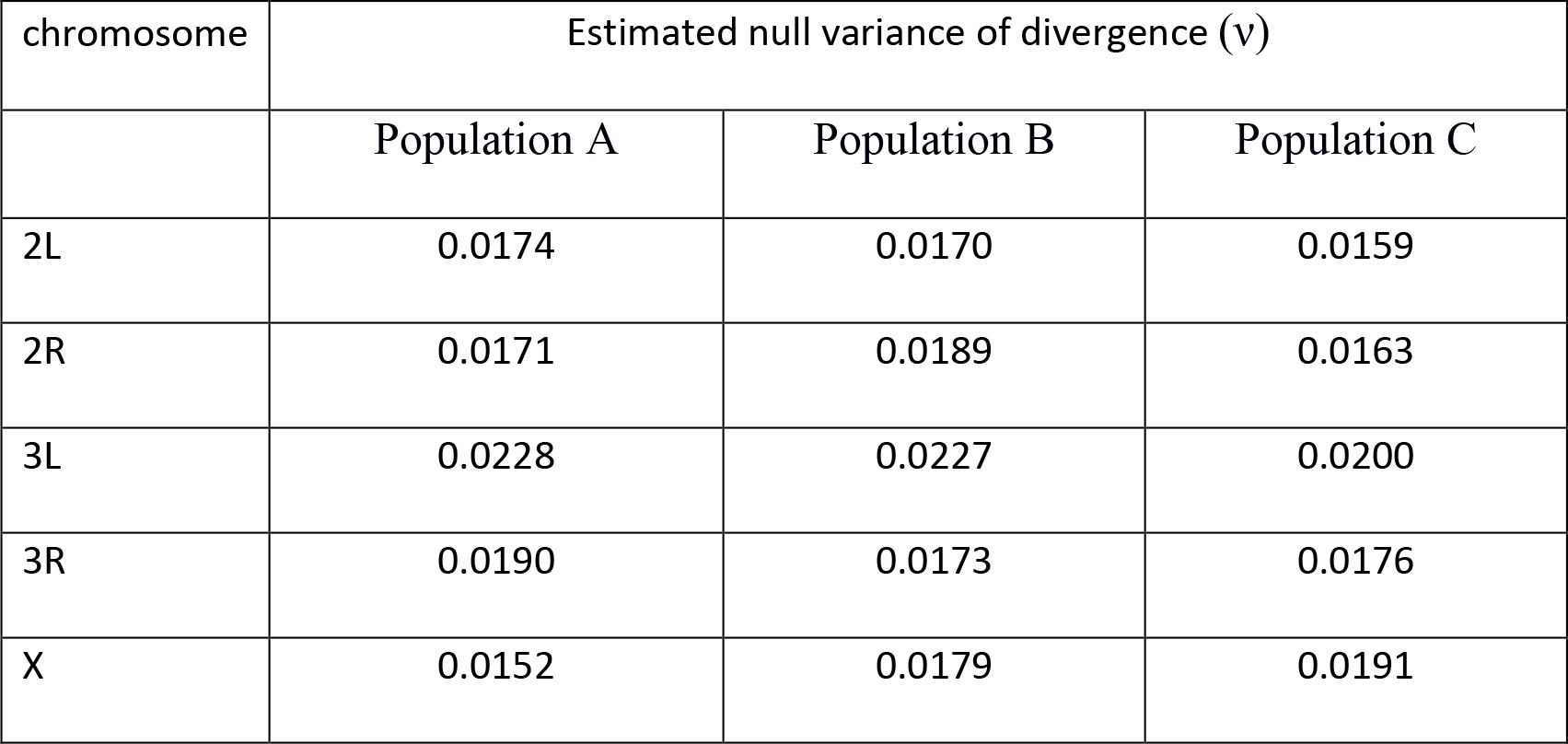
The estimated null variance for each chromosome in each replicate population.

| chromosome | Estimated null variance of divergence (v) |  |  |
| --- | --- | --- | --- |
|  | Population A | Population B | Population C |
| 2L | 0.0174 | 0.0170 | 0.0159 |
| 2R | 0.0171 | 0.0189 | 0.0163 |
| 3L | 0.0228 | 0.0227 | 0.0200 |
| 3R | 0.0190 | 0.0173 | 0.0176 |
| X | 0.0152 | 0.0179 | 0.0191 |

Genomic regions containing LRT-1 significant SNPs exhibit elevated geographical variation (high F_st_) in the MACHADO *et al.* (2016) natural population survey (Table 3). With our filters, the genome-wide mean F_st_ is 0.0150 across the 8 populations (n = 2,340,197 SNPs). The mean is nearly the same if we focus on windows around the 281,917 SNPs of the present experiment that could be ascertained in the r2.01a genome build (F_st_ = 0.0151, n = 330,597 SNPs). Among LRT-1 significant SNPs, we were able to ascertain 119 of 138 loci for subsequent F_st_ calculation.

**Table 3.** The F_st_ estimates for windows around selected sites are compared to the genome wide distribution. F_.025_, F_.975_, and F_.999_ refer to distribution percentiles from sample number of loci (with matching window sizes) from all ascertained loci.

| *window size*<br>*(bp)* | Number of<br>SNPs | $F_{st}$ | $F_{.025}$ | $F_{.975}$ | $F_{.999}$ |
| --- | --- | --- | --- | --- | --- |
| 200 | 332 | **0.0246** | 0.0107 | 0.0191 | 0.0218 |
| 500 | 718 | **0.0242** | 0.0116 | 0.0180 | 0.0200 |
| 1000 | 1246 | **0.0217** | 0.0123 | 0.0177 | 0.0192 |
| 2000 | 2446 | **0.0204** | 0.0126 | 0.0169 | 0.0189 |

Mean F_st_ around selected sites declined with window size (Table 3), but was always much greater for selected loci than for the background genome (45-65% inflation). The F_st_ estimates for selected loci always exceeded even the highest values obtained by resampling (thus p < 0.001 for all window sizes).

### Simulation results

We first performed neutral simulations with N_m_ = N_f_ = 220 following the empirical estimate of N_e_ = 440.9 reported above. As predicted by eqs (1)–(2), the mean *v* across each autosome of each simulated population is very close to the average from Table 2 (0.0185). The LRT distributions are slightly inflated relative to the predicted chi-square distributions (mean LRT-1 = 1.06 instead of 1.0, mean LRT-2= 2.09 instead of 2.0), which suggests that chi-square p-values might be marginally anti-conservative. However, not a single LRT-1 value from any of the 1000 simulation experiments exceeded the permutation threshold (47.0) from the real data. These neutral simulations also establish the null distribution for the CMH tests. We extracted the largest CMH value from each simulation replicate and found the 95^th^ percentile of this list of maxima: 71.5. Returning to the real data and imposing the 71.5 threshold on SNP-specific CMH values, we find that 402 tests exceed 71.5 (Supplemental Table S8) which is ≈3 times the number from LRT-1. All but one of the LRT-1 significant SNPs are within the CMH set.

For simulations with selection, Figure 4 contrasts the multiplicative and truncation selection models over a range of parameter sets in which 30 loci determine fitness (number based on observed results). In these simulations, the selected loci are uniformly positioned over each chromosome. The initial frequencies at selected sites are taken from the 30 ‘selected’ loci in the real data with the specific allele frequency assigned to each SNP randomized with each simulation replicate. The direction of allelic effect was random except when the minor allele was less than 20%. In this situation, the minor allele increased fitness (as in the real data, Supplemental Table S6). The average allele frequency change (x-axis) increases with the strength of selection by either model, but the magnitude of stochastic change generated by linked selection is much greater for multiplicative selection than truncation selection. Here, stochastic change at neutral loci (drift and draft combined) is measured by *v* on the y-axis. In the real experiment, the average estimated allele frequency change at the 30 ‘selected loci’ was about 0.36. At this point in Figure 4, *v* ≈ 0.0185 with for truncation selection but is much greater with multiplicative selection (*v* ≈ 0.032).

**Figure 4.**
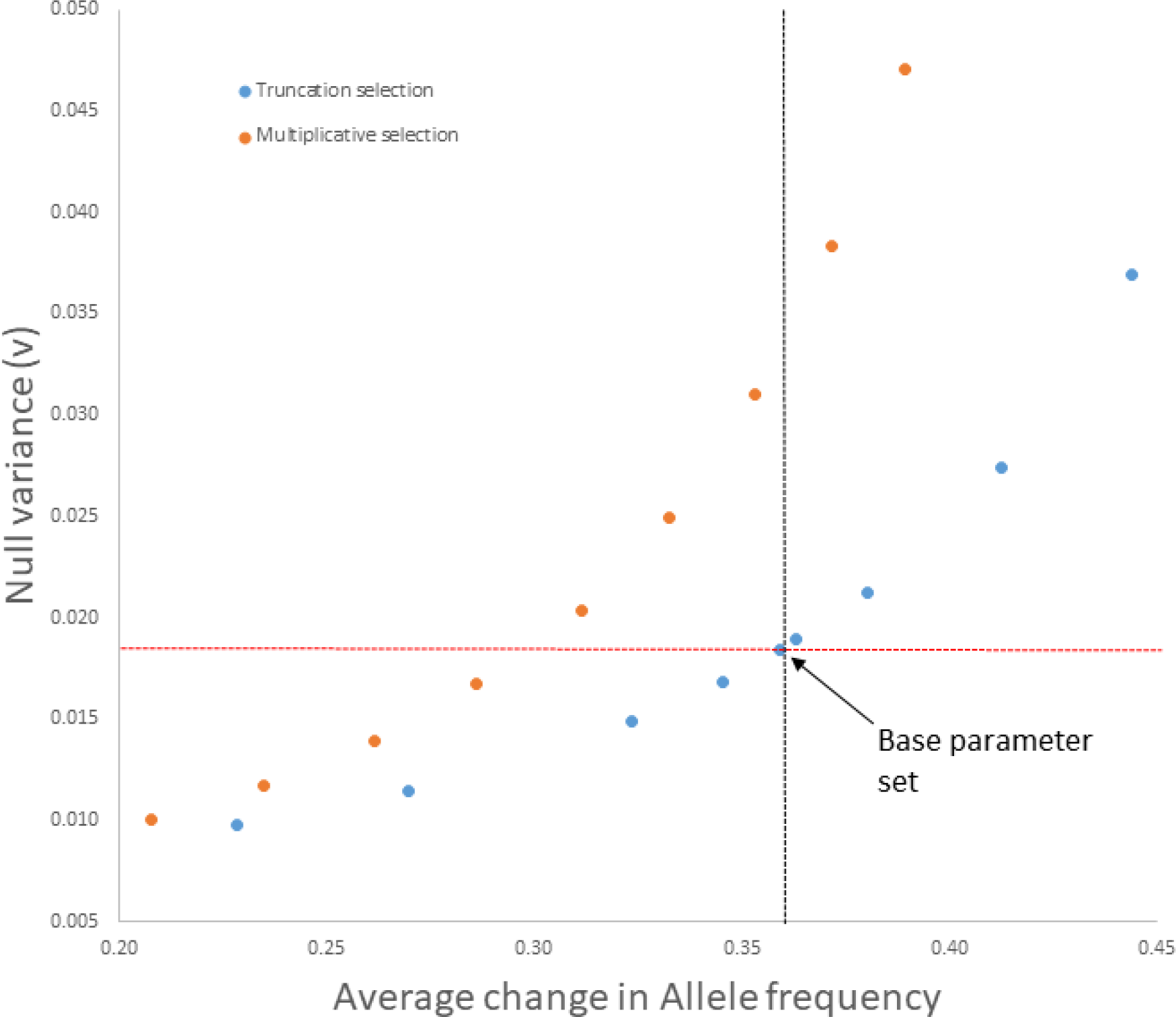
The null variance (mean v across simulation replicates) is reported for varying strengths of truncation selection (blue) and multiplicative selection (orange). The vertical and horizontal lines indicate the mean null variance (red) and mean allele frequency change (grey) from the real experiment. The fractions selected for truncation are 0.8, 0.75, 0.68, 0.65, 0.63, 0.625, 0.6, 0.55, and 0.5, respectively (left to right in graph). The selection coefficients for multiplicative fitness were 0.07, 0.08, 0.09, 0.1, 0.11, 0.12, 0.13, 0.14, and 0.15, respectively. Number of zygotes (pre-selection) was 1600 in all cases. The base parameter set is indicated by arrow.

From the Figure 4 simulations, we chose the truncation model with 63% selected as our “base parameter set” given the match to observed allele frequency change and mean *v*. it also matches the real experiment with respect to the total number of significant LRT-1 tests per simulation replicate (Supplemental Figure S3) as well as the genomic location of significant tests. Specifically, there is aggregation of strong signal in regions of low recombination, illustrated with four randomly chosen replicates in Figure 5 (compare with 3L in Figure 1). Unexpectedly, the 30-locus model generates a large genome-wide correlation of divergence within genomic windows between replicate populations using the Window rank correlation test, which matches the association observed in the data (see shaded distribution in Figure 3). The simulations also reiterate the observed window correlation even if low recombination regions are suppressed (Supplemental Figure S2). Finally, the parameter set with 1600 zygotes and 63% selected yields an adult population size close to our estimated value from the experiment (ca 1000).

**Figure 5.**
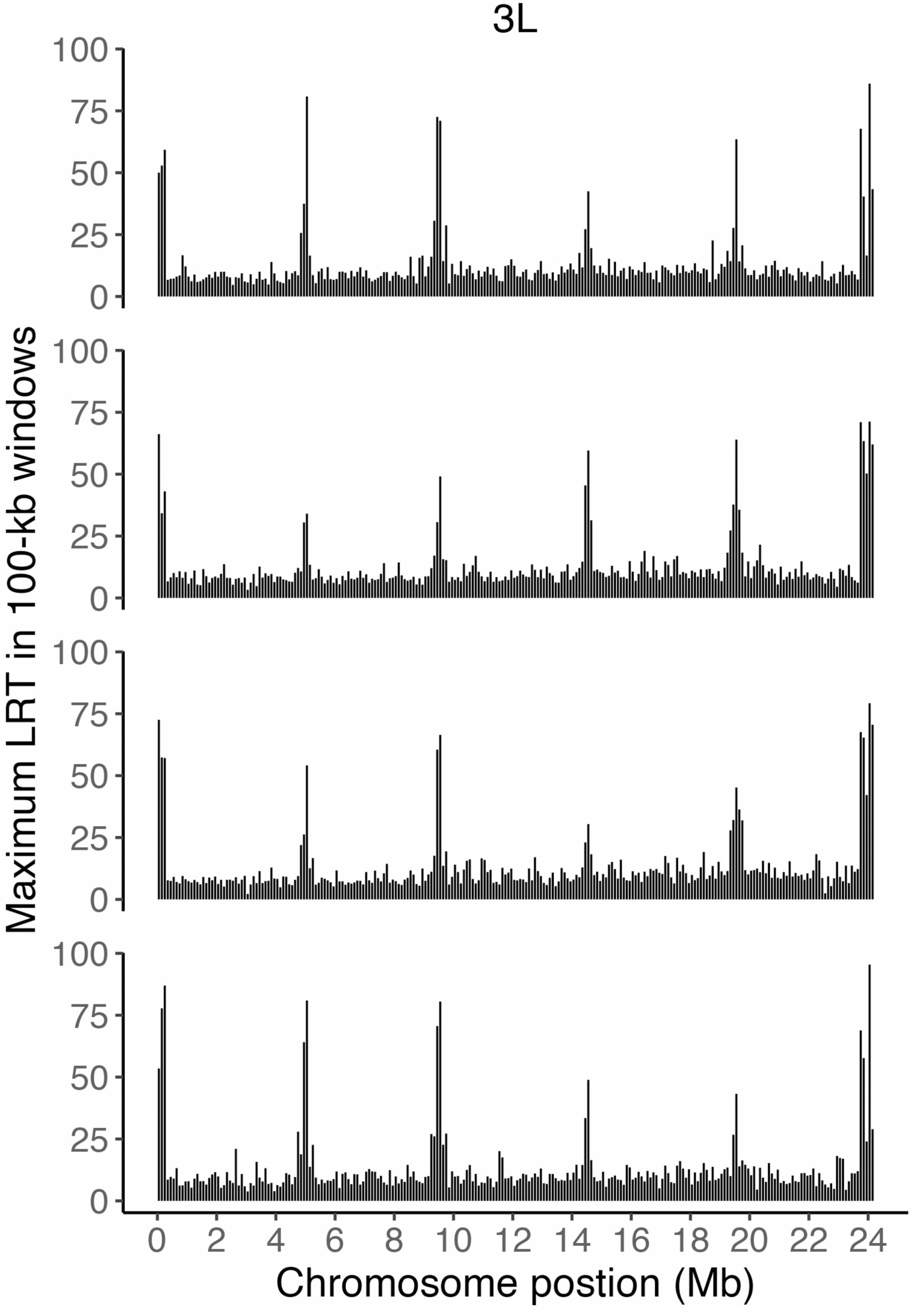
The LRT test is applied to simulated data using the base parameter set (parameters reported in text). The maximum LRT test value for parallel selection per 100kb window along chromosome 3L. Each row represents a different simulation.

Figure 6 compares LRT-1 and CMH tests for the base parameter set using the thresholds from the real experiment (47.0 for LRT-1, 71.5 for CMH). There are over 2 times as many CMH significant SNPs as LRT-1 significant, mirroring the results from the real data (Supplemental Table S8). We can classify each SNP as at a causal site (leftmost group) or as a hitch hiker. The latter is subdivided into three outcomes - within 10kb of selected SNP in a normally recombining region, over 10kb distant from a selected SNP in such a region, or within a low recombination region that contains a selected locus. LRT-1 is considerably more precise in identifying causal SNPs (Figure 6A): 15.2% of LRT-1 tests are at causal SNPs versus 8.6% for CHM. Both tests produce an abundance of significant results in low recombination regions under linked selection (rightmost grouping in Figure 6), but distant hitch-hikers in normal recombination regions are more likely to appear as significant for CMH.

**Figure 6.**
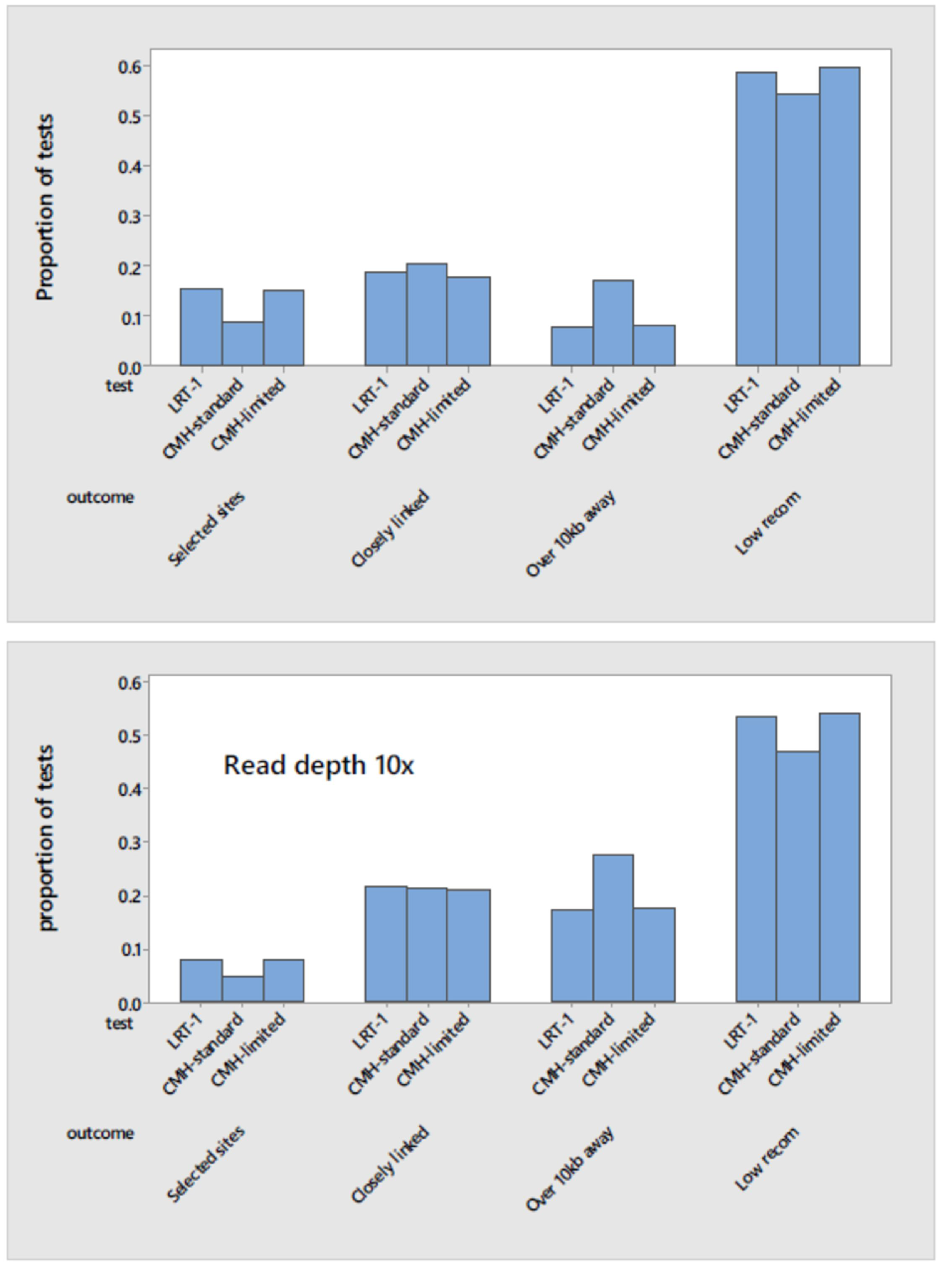
The comparison between LRT-1 and CMH tests is given for the base parameter set using the observed read depths (top) or the read depths increased 10x. Tests are classified as at a fitness determining locus, closely linked (within 10kb) in a normal recombination region, distantly linked in a normal recombination region, or linked but within a low recombination region.

Much of the CMH/LRT-1 difference owes to the fact that CMH is more permissive under the conditions of this experiment (number of populations, read depths, etc.). The CMH-limited category reduces the number of CMH significant tests to the same number as for LRT-1 by inflating the CHM threshold above 71.5 on a replicate specific basis (Figure 6). This greatly reduces the difference between testing methods - precision in identifying causal sites is only slightly higher for LRT-1 than CMH-limited (within 10%) and many of the CMH significant tests that are far from selected sites are erased. We also considered the effect of increasing read depths because our experiment had relatively low depths, 5-10% of the number of chromosomes sampled into each bulk except for C7 (25%). We performed simulations of the base parameter set but with read depths increased 10 fold (Figure 6B). To establish the appropriate significance threshold for CMH, we repeated the neutral simulations with elevated read depths (new threshold = 437.0). As expected, the number of significant tests increases for both methods (2.5× increase for LRT-1, 1.9× for CMH). However, the fraction of tests at selected sites declines. Nearly all selected sites are identified by both methods (average 29 of 30) but the great majority of new significant tests with increased read depth are hitch-hikers.

To evaluate the effects of LD on genome-wide patterns, we eliminated LD and simulated the base parameter set with all else unchanged (including limited recombination within centro/telomeric regions). The results are profoundly different. There is no effect on the mean (absolute) allele frequency change at selected sites, but the number of significant LRT-1 tests is dramatically reduced (from an average of 149 to 23.6). However, every single significant LRT-1 test across 1000 simulation replicates occurred at a causal site, illustrating the profound effect of hitch-hiking even in a species with relatively low LD.

We next manipulated the initial distribution of allele frequencies at selected loci to address the question that emerges from Figure 2A: Does the intermediate frequency of significant variants from the experiment imply that intermediate frequency variants are the primary targets of selection in the laboratory environment? We conducted simulations using the base parameter set except with initial allele frequencies of selected loci sampled from the genome-wide distribution (blue bars in Figure 2A; Supplemental Table S6.E). This change actually increases the average number of significant LRT-1 tests (from 149 to 182), but the number of significant tests at causal SNPs declines. The increase in significant hitch-hikers suggests that linked selection has a more pronounced effect when the favored allele is initially uncommon.

The broken lines (green) in Figure 2A illustrate the initial AFS of LRT-1 significant tests from these simulations. There is a “pull to the middle” - most selected sites have a MAF < 0.1, but those with higher initial frequencies are more likely to yield high LRT-1 (Figure 2A). Despite this ascertainment effect (among selected sites, those with intermediate frequency are more likely to be detected), the real data contain an excess of SNPs with MAF > 0.3 that yield significant tests (orange versus green in Figure 3). The pull to the middle in simulations with selection from the background AFS reflects a biologically important feature - positively selected loci with higher MAF are generating greater variance in fitness when the population experiences the novel laboratory environment.

## Discussion

In this E&R experiment, we observed parallel changes in allele frequency at over a 100 SNPs, clustered into about 30 distinct loci across the genome. The allele frequency spectrum (AFS) of these putatively selected sites is strongly biased toward intermediate allele frequencies. The source natural population clearly harbors abundant standing variation to allow rapid adaptation in a novel environment. Accumulating examples of rapid phenotypic and genomic evolution and observations that allele frequencies can vary cyclically in natural Drosophila populations (BERGLAND *et al.* 2014) have led to the suggestion that many of the foundational principles of molecular population genetics, based on neutral and nearly-neutral theory, might require revision (MESSER *et al.* 2016; HERMISSON AND PENNINGS 2017). In this experiment, we could not find a neutral model that accurately predicts the evolution of the typical SNP. Adjusting N_e_ allows the neutral model to predict the average change in frequency (mean v), but not the elevated variability of change (Table 2) or the covariance of change in genomic windows across independent replicated populations (Figure 3). Below, we discuss these results in relation to the maintenance of polymorphisms in nature and the paradox of excessive significance in E&R studies.

### The maintenance of polymorphism

We hypothesize that our selected polymorphisms (Table S6) are enriched for loci under balancing selection in nature (BERGLAND *et al.* 2014; CHARLESWORTH 2015). This hypothesis is based on two features of the results. First, the AFS of LRT-1 significant SNPs is strikingly different from the genomic background. The overall AFS (blue bars in Figure 2A) exhibits the expected preponderance of rare alleles (MORIYAMA AND POWELL 1996; PRZEWORSKI *et al.* 2001), consistent with the idea that most polymorphisms are neutral or nearly neutral (WRIGHT 1931; OHTA 1976). However, the polymorphisms that responded to selection are intermediate in frequency (orange bars in Figure 2A). In molecular population genetics, the most common AFS-based test for selection is Tajima’s D (TAJIMA 1989), which has an expected value of zero under the equilibrium neutral model. At the gene level, Tajima’s D > 2.0 is considered significantly positive (assuming that large number of alleles are sequenced) and typically interpreted as evidence for balancing selection. For our selected SNPs, Tajima’s D = 4.88 (using the median Ancestral read depth of 324 for n), hugely inflated relative to the neutral expectation and also in comparison to the genomic background value.

Balancing selection can result in a stable minor allele frequency less than 10% (in the lowest category of Figure 2A), but deleterious variants are unlikely to segregate at intermediate frequencies (higher categories in Figure 2A). Neutral variants may occasionally drift to intermediate frequencies, but the overall AFS of our LRT-1 SNPs is not consistent with neutrality. Ascertainment is an important consideration here: Does the excess of intermediate frequency polymorphisms result simply because change at these loci is easier to detect? For the great majority of polymorphisms, the answer is clearly no. The average magnitude of change is as large for rare alleles as for common (Figure 2B) because the angular transformation effectively normalizes the effects of genetic drift and experimental sampling on allele frequency change (FISHER AND FORD 1947; WALSH AND LYNCH 2018). However, the transformation does not eliminate the dependency for selection-driven change because the variance in fitness is maximal at intermediate allele frequency. The difference between blue and green in Figure 2 illustrates that we are more likely to detect intermediate frequency alleles because they experience greater change with the same strength of selection. However, even after correcting for this effect, the real data still yield an excess of significant SNPs with MAF > 0.3 (orange versus green in Figure 2A).

Another recent E&R experiment on *D. simulans* provides an empirical demonstration that evolution of rare alleles to intermediate frequencies is detectable, if that is the basis of the response (see Fig 2 of BARGHI *et al.* (2017)). In that study, loci that evolved significantly generally had MAF < 0.2 and became more intermediate in frequency during laboratory adaptation. There are a number of differences between the studies that could explain the differing outcomes. For example, the BARGHI *et al.* (2017) founders were derived from isofemale lines, whereas the founders in our experiment were the offspring of wild-caught females. Regardless of the reason, the results of BARGHI et al. (2017) indicate that rare, favorable alleles have detectable signal. If we run our simulator using the base parameter set, except replacing the observed AFS with an initial frequency of 0.02 for all favorable alleles, the number of significant LRT-1 tests actually increases. Mean allele frequency at selected SNPs is reduced (from 0.36 to 0.23), as is the number of significant LRT-1 tests at those selected SNPs. However, the inflation of hitch-hiking, particularly beyond 10kb more than compensates.

A second observation, relevant to natural selection on our significant SNPs, is that the genomic regions harboring these SNPs exhibit elevated differentiation among natural populations of *D. simulans* in Eastern North America (Table 3; MACHADO *et al.* (2016)). This pattern suggests that alternative alleles at our significant SNPs might be responsive to environmental heterogeneity (LEWONTIN AND KRAKAUER 1973; BEAUMONT AND NICHOLS 1996). We have emphasized that the lab environment is a novel selective challenge to wild *D. simulans*, but it is also relatively constant and homogeneous. With heterogeneity in the natural environment (e.g., seasonal variation in temperature or spatial structure in resource availability), a ‘multi-niche polymorphism’ (LEVENE 1953) can result if genotypes vary in their environmental optima. Such polymorphisms will not remain stable if environmental heterogeneity is eliminated. We cannot predict which allele would increase in captivity without detailed information about genotype-specific tolerances. However, it is likely that one genotype will, by chance, match the lab environment better than alternatives, resulting in the eventual loss of the latter. Frequency-dependent selection arising from competitive or social interactions (ANTONOVICS AND KAREIVA 1988) can also maintain polymorphism that the lab environment would remove or reduce.

### Hitch-hiking and excessive significance

NUZHDIN AND TURNER (2013) argue that the large number of significant tests in recent E&R experiments, e.g. (TURNER *et al.* 2011; OROZCO-TER WENGEL *et al.* 2012; TURNER AND MILLER 2012), must be due to over-estimating the number of loci under selection. Because a limited number of haplotypes are sampled to initiate E&R experiments, non-random associations between loci can occur between loci far apart in the genome. This sampling effect, combined with traditional hitch-hiking, could produce large changes in allele frequency at many sites that are not the direct targets of selection. For the present experiment, we established each replicate population with a distinct sampling of genotypes from nature, which should reduce the scope for haplotype-sampled LD to generate false positives. The finite number of haplotypes that survive in each experimental population will yield idiosyncratic associations between distant SNPs, but these associations should be population specific, and thus less likely to generate consistent *parallel* changes across replicates.

While founding experimental populations with distinct samples from the natural population may reduce sampling LD, it will not eliminate genuine long distance LD, that present in the natural population. Our treatment of LD in the simulations is based on the inbred line sequencing of SIGNOR *et al.* (2018) which does reveal non-trivial levels of LD, particularly in low recombination regions (Supplemental Tables S3-S4, Figure S1). The simulations indicate that hitch-hiking based on this level of LD still has a pervasive effect (Figures 3-5). Perhaps most surprising is that selection on only 30 loci generates a genome-wide positive association between locus specific responses across independent replicated populations (Figures 3, S2). It is true that the SIGNOR *et al.* (2018) data, on which we calibrate our simulations, is itself a finite sample of ca. 170 alleles. It might thus have its own collection of sampling induced LD values. However, artificial LD is most likely to occur between rare variants that are accidentally captured in the same line or lines. The hitch-hiking in our simulations is driven by associations between intermediate frequency alleles where the “minor haplotype” is sampled into many sequenced lines (and correspondingly into each of the replicated populations of our simulations). These estimates are far more robust. An important corollary is that, among our significant SNPs (Supplemental Table S6), we cannot distinguish the actual targets of selection from hitch-hikers. While this is a clear limitation of the experiment, the intermediate frequency result (Figure 2A) is not undermined by hitch-hiking. Change at a hitch-hiking locus will only match change at the selected locus if allelic association is maximal (and remains unbroken by recombination) and allele frequencies are similar at the two loci (r^2^ near 1).

### Significance testing

The number of significant tests (138 for LRT-1) is reduced by orders of magnitude from previous E&R studies owing to differences in both experimental design and statistical analysis. The independent founding of experimental populations (described above) is one factor, but minimum depth thresholds also caused many of our SNPs to drop out of the set tested for selection. The simulation results of Figure 6B suggest that increased read depth would increase the number of significant tests, although the predicted inflation is caused overwhelming by the inclusion of additional hitch-hikers rather than novel selected sites. Differing testing methods also contribute - we obtain two to three times as many significant tests for CMH than LRT-1 when applied to the same dataset (either real or simulated). CMH and LRT-1 assimilate read counts in different ways. The contingency table method (CMH) should work best when each sequenced read is an independently sampled allele from the relevant population (Ancestral or Descendant in this case). This is not a requirement for LRT-1, but the independence assumption likely holds fairly well for this experiment given that read counts were far below the number of alleles in each population. Thus, the difference is most likely caused by the different ways that significance thresholds are determined, neutral simulation for CMH, permutation for LRT-1, with the latter more conservative.

The four tests in Table 1 are intended to provide complimentary information about the genome wide response to selection. In this experiment, the Extreme windows test was redundant with LRT-1 and only one SNP was genome-wide significant for heterogeneous selection (LRT-2). This is surprising, because even if environmental conditions were exactly the same across replicates, stochastic loss of alleles could generate significant LRT-2 tests. Rare alleles in the natural population might be randomly excluded from one or more of the experimental replicates. If the rare allele is favorable, it will increase where present but not where absent, yielding a heterogeneous response across replicates. Direct inspection of the full panel of SNPs (Supplemental Table S1) indicates that this very rarely happened. The minor allele was absent from at least one replicate population at 13,856 SNPs (ca. 5% of cases). At these SNPs, polymorphic populations showed very low average change. There were a small number of SNPs with substantial change, but in these the response was limited to a single replicate and magnitude of allele frequency change (0.10-0.15) was much lower than at LRT-1 significant SNPs (average change 0.35). These observations support the claim that adaptive evolution in this experiment mainly involved alleles that segregated at intermediate frequency in the ancestral wild population.

A biological reason for fewer significant SNPs here relative to E&R experiments on *D. melanogaster* (BURKE *et al.* 2010; TURNER *et al.* 2011; OROZCO-TERWENGEL *et al.* 2012) is that inversions are rare in *D. simulans* (Lemeunier and Aulard 1992) and the genome-wide recombination rate is 30% higher (True et al. 1996). BARGHI *et al.* (2017) compared genomic change in *D. melanogaster* and *D. simulans* after ~60 generations of hot laboratory conditions and found differing patterns between the two species. *D. simulans* had fewer candidate SNPs, and the regions of the genome implicated in response to selection were narrower and more distinct. Strikingly, almost all of chromosome arm 3R in *D. melanogaster* (which contains several overlapping segregating inversions) exhibited a pattern consistent with selection. In *D. simulans*, which lacks similar inversions on 3R, several narrow, distinct regions on this chromosome arm exhibited such a pattern. The authors attributed many of these differences in the frequency of segregating inversions and in centromeric recombination suppression.

### Prospects

Predicting the evolutionary response to a changing environment, and its genetic basis, is one of the major goals of modern evolutionary genetics. We combine a genomic sequencing experiment with a simulation model tailored as closely as possible to the relevant features of that experiment. The simulation demonstrates that a strong response at a limited number of loci can generate a genome-wide response involving thousands of polymorphic sites through hitch-hiking, consistent with the observations of the experiment. A more ambitious application would be to use the simulator for formal inference, to identify selected SNPs and the strength of selection on each. In principle, the simulator could be employed for Approximate Bayesian Computation (BEAUMONT *et al.* 2002) or as a trainer for Supervised Machine Learning (SCHRIDER AND KERN 2018), but Figure 4 suggests a primary question. What should we attempt to estimate? Allele frequency change under multi-locus selection depends on how locus-specific effects combine to determine survival and reproduction. We found a generally better fit to the data with truncation rather than multiplicative selection; the latter inducing an excessive amount of draft to achieve the same magnitude of allele frequency change at selected loci. This observation is consistent with theoretical studies of ‘substitution load’ from over 50 years ago (HALDANE 1957; SVED *et al.* 1967; KIMURA AND MARUYAMA 1969; WALLACE 1970). While our base parameter set with truncation selection reiterates the major observations of the study, we do not think it fully accurate. The conundrum is that while both models (truncation and multiplicative selection) offer parameters to estimate, interpretation of estimates is problematic if neither model describes the genotype-to-fitness mapping.

The most important biological conclusions from this experiment follow from the AFS of selected loci (Fig 2). The founding of our experimental populations may have been a critical determinant of the intermediate frequency result. Our ancestral populations were only one generation removed from nature, did not experience population bottlenecks, and did not undergo multiple generations of laboratory adaptation or inbreeding prior to the start of the experiment. Lab adaptation, inbreeding, or population contraction in the founders would likely have changed the starting frequencies. This aspect of experimental design not only affects patterns of evolution in an E&R experiment, but also inferences regarding the selective forces acting on these loci in nature.

## Acknowledgments

We thank S. Signor, H. Machado, T. O’Connor, R. Unckless, S. Macdonald, D. Houle, and J. Sztepanacz for input on the project and/or manuscript. We thank Christopher Souders and Lauren Reynolds for lab work and the KU ACF for computing support.

## Supplemental Materials

**Table S1. The full set of responses at 291,272 SNPs.**

**Table S2. SNPs significant for LRT-1 where a polymorphisms was also observed in the Machado (columns E-H) or Signor (columns I-L) datasets. Mis-matches are color coded.**

**Table S3. The hash tables for two locus haplotype (gamete) frequencies are reported as a function of allele frequencies (minor allele at both loci) and the distance between loci. Separate tables are given for autosomes and the X chromosome. The calculation procedures are described on front sheet.**

**Table S4. A summary of LD estimates (measured as r^2^) from with distances ranging from 1 bp to 300,000bp. Inter-SNP distances are binned into six categories: 0 if distance less than 10bp, 1 if less than 100bp, 2 if less than 1kb, 3 if less than 10kb, 4 if less than 100kb, and 5 if greater than 100kb. Each window contains the relevant contrasts within that part of the genome.**

**Table S5. The crossovers location distribution used for simulation based on data from D. melanogaster from Comeron et al. (2012). Rates are reduced rates near telomeres and centromeres.**

**Table S6. The 138 polymorphisms with an LRT-1 value (test for parallel selection) that exceed the permutation threshold of 47.0. The last column (Locus30?) identifies the subset of SNPs that are most significant in each locus (as 1).**

**Table S7. The maximum LRT-1 value (genome-wide) is reported for each of 1000 permutations of the data. The threshold is based on 95th percentile of this distribution.**

**Table S8. All SNPs that were significant by either LRT-1 or CMH. The rankings are within the set of LRT-1 (column G) or CMH (column H). ns = non-significant**

**Table S9. The summary of results in 25 SNP genomic windows.**

**Supplemental Figure S1. The average value for LD measured as r is reported as a function of chromosomal location with the “Sz” D. simulans lines published by Signor et al. (2018). r is for pairs of loci separated by 100-300kb and where the minor allele at each locus is greater than 10%. The red line is the centromere for chromosomes 2 and 3.**

**Supplemental Figure S2. The average Spearman correlation is indicated relative to the distribution obtained by permutation of windows (white bars) and from simulations with selection on 30 loci (filled bars). These correlations were calculated after excluding top 21 most diverged windows and windows in low-recombination regions surrounding centromeres and telomeres.**

**Supplemental Figure S3. The distribution of significant LRT-1 tests per experiment is given for 1000 simulations of the base parameter set. The vertical blue lines denotes the number observed in the real experiment.**

**Supplemental.File.1.tar.gz :: The python programs LD.analysis1.py, LD.analysis2.py, LD.analysis3.py used for Table S4. The c program to simulate the experiment.**

## Literature Cited

Antonovics, J., and P. Kareiva, 1988 Frequency-dependent selection and competition - empirical approaches. Phil. Trans. Roy. Soc. Lond. B 319: 601–613.

Barghi, N., R. Tobler, V. Nolte and C. Schlotterer, 2017 Drosophila simulans: A Species with Improved Resolution in Evolve and Resequence Studies. G3 (Bethesda) 7: 2337–2343.

Barrick, J. E., D. S. Yu, S. H. Yoon, H. Jeong, T. K. Oh et al., 2009 Genome evolution and adaptation in a long-term experiment with Escherichia coli. Nature 461: 1243–U1274.

Barton, N. H., 1995 Linkage and the Limits to Natural-Selection. Genetics 140: 821–841.

Beaumont, M. A., and R. A. Nichols, 1996 Evaluating loci for use in the genetic analysis of population structure. Proceedings Of the Royal Society Of London Series B-Biological Sciences 263: 1619–1626.

Beaumont, M. A., W. Zhang and D. J. Balding, 2002 Approximate Bayesian Computation in Population Genetics. Genetics 162: 2025–2035.

Beissinger, T. M., C. N. Hirsch, B. Vaillancourt, S. Deshpande, K. Barry et al., 2014 A Genome-Wide Scan for Evidence of Selection in a Maize Population Under Long-Term Artificial Selection for Ear Number. Genetics 196: 829–840.

Beissinger, T. M., G. J. M. Rosa, S. M. Kaeppler, D. Gianola and N. de Leon, 2015 Defining window-boundaries for genomic analyses using smoothing spline techniques. Genetics Selection Evolution 47: 30.

Bergland, A. O., E. L. Behrman, K. R. O’Brien, P. S. Schmidt and D. A. Petrov, 2014 Genomic Evidence of Rapid and Stable Adaptive Oscillations over Seasonal Time Scales in Drosophila. PLOS Genetics 10: e1004775.

Burke, M. K., J. P. Dunham, P. Shahrestani, K. R. Thornton, M. R. Rose et al., 2010 Genome-wide analysis of a long-term evolution experiment with Drosophila. Nature 467: 587–U111.

Charlesworth, B., 2009 Effective population size and patterns of molecular evolution and variation. Nat Rev Genet 10: 195–205.

Charlesworth, B., 2015 Causes of natural variation in fitness: Evidence from studies of Drosophila populations. Proceedings of the National Academy of Sciences 112: 1662–1669.

Comeron, J. M., R. Ratnappan and S. Bailin, 2012 The Many Landscapes of Recombination in Drosophila melanogaster. PLOS Genetics 8: e1002905.

Conover, D. O., and S. B. Munch, 2002 Sustaining fisheries yields over evolutionary time scales. Science 297: 94–96.

Corbett-Detig, R. B., and D. L. Hartl, 2012 Population genomics of inversion polymorphisms in Drosophila melanogaster. PLoS Genet 8: e1003056.

Daborn, P. J., J. L. Yen, M. R. Bogwitz, G. Le Goff, E. Feil et al., 2002 A single p450 allele associated with insecticide resistance in Drosophila. Science 297: 2253–2256.

Darimont, C. T., S. M. Carlson, M. T. Kinnison, P. C. Paquet, T. E. Reimchen et al., 2009 Human predators outpace other agents of trait change in the wild. Proceedings of the National Academy of Sciences of the United States of America 106: 952–954.

Ellner, S. P., M. A. Geber and N. G. Hairston, Jr., 2011 Does rapid evolution matter? Measuring the rate of contemporary evolution and its impacts on ecological dynamics. Ecol Lett 14: 603–614.

Ellner, S. P., N. G. Hairston, Jr., C. M. Kearns and D. Babai, 1999 The roles of fluctuating selection and long-term diapause in microevolution of diapause timing in a freshwater copepod. Evolution 53: 111–122.

Fisher, R. A., and E. B. Ford, 1947 The spread of a gene in natural conditions in a colony of the moth Panaxia dominula. Heredity 1: 143–174.

Ford, E. B., 1964 Ecological Genetics. Methuen and Company, London, UK.

Franks, S. J., and A. E. Weis, 2008 A change in climate causes rapid evolution of multiple life-history traits and their interactions in an annual plant. Journal of Evolutionary Biology 21: 1321–1334.

Franssen, S. U., V. Nolte, R. Tobler and C. Schlötterer, 2015 Patterns of Linkage Disequilibrium and Long Range Hitchhiking in Evolving Experimental Drosophila melanogaster Populations. Molecular Biology and Evolution 32: 495–509.

Ghalambor, C. K., K. Hoke, E. Ruell, E. K. Fisher, D. N. reznick et al., 2015 Non-adaptive plasticity and rapid adaptive evolution of the guppy transcriptome. Nature 525: 372–375.

Gillespie, J. H., 1991 The Causes of Molecular Evolution. Oxford University Press, Oxford.

Gillespie, J. H., 2001 IS THE POPULATION SIZE OF A SPECIES RELEVANT TO ITS EVOLUTION? Evolution 55: 2161–2169.

Grant, P. R., 1999 Ecology and Evoution of Darwin&#x2019;s Finches. Princeton University Press, Princeton, N.J.

Hairston, N. G., S. P. Ellner, M. A. Geber, T. Yoshida and J. A. Fox, 2005 Rapid evolution and the convergence of ecological and evolutionary time. Ecology Letters 8: 1114–1127.

Hairston, N. G., and W. E. Walton, 1986 Rapid evolution of a life history trait. Proc Natl Acad Sci U S A 83: 4831–4833.

Haldane, J. B. S., 1957 The cost of natural selection. J. Genet. 55: 511–524.

Hendry, A. P., and M. T. Kinnison, 1999 Perspective: The pace of modern life: Measuring rates of contemporary microevolution. Evolution 53: 1637–1653.

Hermisson, J., and P. S. Pennings, 2017 Soft sweeps and beyond: understanding the patterns and probabilities of selection footprints under rapid adaptation. Methods in Ecology and Evolution 8: 700–716.

Hu, T. T., M. B. Eisen, K. R. Thornton and P. Andolfatto, 2013 A second-generation assembly of the Drosophila simulans genome provides new insights into patterns of lineage-specific divergence. Genome Research 23: 89–98.

Huang, Y., and A. F. Agrawal, 2016 Experimental evolution of gene expression and plasticity in alternative selective regimes. PLoS Genet 12: e1006336.

Huang, Y., S. I. Wright and A. F. Agrawal, 2014 Genome-wide patterns of genetic variation within and among alternative selective regimes. PLoS Genet 10: e1004527.

Jain, K., and W. Stephan, 2017 Rapid adaptation of a polygenic trait after a sudden environmental shift. Genetics 206: 389–406.

Johnston, R. F., and R. K. Selander, 1964 House sparrows: rapid evolution of races in North America. Science 144: .548–550.

Kang, L., D. D. Aggarwal, E. Rashkovetsky, A. B. Korol and P. Michalak, 2016 Rapid genomic changes in Drosophila melanogaster adapting to desiccation stress in an experimental evolution system. BMC Genomics 17: 233.

Kapun, M., H. van Schalkwyk, B. McAllister, T. Flatt and C. Schlotterer, 2014 Inference of chromosomal inversion dynamics from Pool-Seq data in natural and laboratory populations of Drosophila melanogaster. Mol Ecol 23: 1813–1827.

Kelly, J. K., B. Koseva and J. P. Mojica, 2013 The Genomic Signal of Partial Sweeps in Mimulus guttatus. Genome Biology and Evolution 5: 1457–1469.

Kettlewell, H. D., 1958. A survey of the frequencies of Biston betularia (L.)(Lep.) and its melanic forms in Great Britain. Heredity 12: 51–72.

Kimura, M., and T. Maruyama, 1969 The substitutional load in a finite population1. Heredity 24: 101.

Koboldt, D. C., K. Chen, T. Wylie, D. E. Larson, M. D. McLellan et al., 2009 VarScan: variant detection in massively parallel sequencing of individual and pooled samples. Bioinformatics 25: 2283–2285.

Kofler, R., and C. Schlötterer, 2014 A Guide for the Design of Evolve and Resequencing Studies. Molecular Biology and Evolution 31: 474–483.

Lang, G. I., D. P. Rice, M. J. Hickman, E. Sodergren, G. M. Weinstock et al., 2013 Pervasive genetic hitchhiking and clonal interference in forty evolving yeast populations. Nature 500:571–574.

Levene, H., 1953 Genetic equilibrium when more than one ecological niche is available. Amer. Natur. 87: 331–333.

Levy, S. F., J. R. Blundell, S. Venkataram, D. A. Petrov, D. S. Fisher et al., 2015 Quantitative evolutionary dynamics using high-resolution lineage tracking. Nature 519: 181–186.

Lewontin, R. C., 1964 THE INTERACTION OF SELECTION AND LINKAGE. I. GENERAL CONSIDERATIONS; HETEROTIC MODELS. Genetics 49: 49–67.

Lewontin, R. C., 1974 The genetic basis of evolutionary change. Columbia University Press, New York, NY.

Lewontin, R. C., and J. Krakauer, 1973 Distribution of gene frequency as a test of the theory of the selective neutrality of polymorphisms. Genetics 74: 175–195.

Li, H., and R. Durbin, 2009 Fast and accurate short read alignment with Burrows-Wheeler transform. Bioinformatics 25: 1754–1760.

Li, H., B. Handsaker, A. Wysoker, T. Fennell, J. Ruan et al., 2009 The Sequence Alignment/Map format and SAMtools. Bioinformatics 25: 2078–2079.

Long, A., G. Liti, A. Luptak and O. Tenaillon, 2015 Elucidating the molecular architecture of adaptation via evolve and resequence experiments. Nature Reviews Genetics 16: 567.

Losos, J. B., K. I. Warheit and T. W. Shoener, 1997 Adaptive differentiation following experimental island colonization in Anolis lizards. Nature 387: 70–73.

Machado, H. E., A. O. Bergland, K. R. O’Brien, E. L. Behrman, P. S. Schmidt et al., 2016 Comparative population genomics of latitudinal variation in Drosophila simulans and Drosophila melanogaster. Molecular Ecology 25: 723–740.

Maynard Smith, J., and J. Haigh, 1974 The hitch-hiking effect of a favourable gene. Genetic research 23: 23–35.

McKenna, A., M. Hanna, E. Banks, A. Sivachenko, K. Cibulskis et al., 2010 The Genome Analysis Toolkit: A MapReduce framework for analyzing next-generation DNA sequencing data. Genome Research 20: 1297–1303.

Messer, P. W., S. P. Ellner and N. G. Hairston, Jr., 2016 Can population genetics adapt to rapid evolution? Trends Genet 32: 408–418.

Michalak, P., L. Kang, P. M. Sarup, M. F. Schou and V. Loeschcke, 2017 Nucleotide diversity inflation as a genome-wide response to experimental lifespan extension in Drosophila melanogaster. BMC Genomics 18: 84.

Monnahan, P. J., and J. K. Kelly, 2017 The Genomic Architecture of Flowering Time Varies Across Space and Time in<em>Mimulus guttatus<em>. Genetics.

Moriyama, E. N., and J. R. Powell, 1996 Intraspecific Nuclear-Dna Variation In Drosophila. Molecular Biology and Evolution 13: 261–277.

Neher, R. A., and B. I. Shraiman, 2011 Genetic Draft and Quasi-Neutrality in Large Facultatively Sexual Populations. Genetics 188: 975–996.

Nuzhdin, S. V., and T. L. Turner, 2013 Promises and limitations of hitchhiking mapping. Current Opinion in Genetics &#x0026; Development 23: 694–699.

Ohta, T., 1976 Role of very slightly deleterious mutations in molecular evolution and polymorphism. Theor. Pop. Biol. 10: 254–275.

Orozco-terWengel, P., M. Kapun, V. Nolte, R. Kofler, T. Flatt et al., 2012 Adaptation of Drosophila to a novel laboratory environment reveals temporally heterogeneous trajectories of selected alleles. Molecular Ecology 21: 4931–4941.

Palumbi, S. R., 2001 The Evolution Explosion: How Humans Cause Rapid Evolutionary Change. W.W. Norton, New York, NY, USA.

Przeworski, M., J. D. Wall and P. Andolfatto, 2001 Recombination and the Frequency Spectrum in Drosophila melanogaster and Drosophila simulans. Molecular Biology and Evolution 18: 291–298.

Remolina, S. C., P. L. Chang, J. Leips, S. V. Nuzhdin and K. A. Hughes, 2012 Genomic basis of aging and life-history evolution in Drosophila melanogaster. Evolution 66: 3390–3403.

Reznick, D. N., F. H. Shaw, F. H. Rodd and R. G. Shaw, 1997 Evaluation of the rate of evolution in natural populations of guppies (Poecilia reticulata). Science 275: 1934–1937.

Rose, M. R., 1984 Laboratory evolution of postponed senescence in Drosophila melanogaster. Evolution 38: 1004–1010.

Schou, M. F., V. Loeschcke, J. Bechsgaard, C. Schlotterer and T. N. Kristensen, 2017 Unexpected high genetic diversity in small populations suggests maintenance by associative overdominance. Mol Ecol.

Schrider, D. R., and A. D. Kern, 2018 Supervised Machine Learning for Population Genetics: A New Paradigm. Trends in Genetics 34: 301–312.

Signor, S. A., F. N. New and S. Nuzhdin, 2018 A Large Panel of Drosophila simulans Reveals an Abundance of Common Variants. Genome Biology and Evolution 10: 189–206.

Slobodkin, L. B., 1980 Growth and Regulation in Animal Populations. Dover, New York, NY.

Stuart, Y. E., T. S. Campbell, P. A. Hohenlohe, R. G. Reynolds, L. J. Revell et al., 2014 Rapid evolution of a native species following invasion by a congener. Science 346: 463–466.

Sukumaran, J., and M. T. Holder, 2011 Ginkgo: spatially-explicit simulator of complex phylogeographic histories. Molecular Ecology Resources 11: 364–369.

Sved, J. A., T. E. Reed and W. F. Bodmer, 1967 The number of balanced populations that can be maintained in a natural population. Genetics 55: 469–481.

Tajima, F., 1989 Statistical Method for Testing the Neutral Mutation Hypothesis by DNA Polymorphism. Genetics 123: 585–595.

Thompson, J. N., 1998 Rapid evolution as an ecological process. Trends in Ecology & Evolution 13: 329–332.

Tobler, R., S. U. Franssen, R. Kofler, P. Orozco-Terwengel, V. Nolte et al., 2014 Massive habitat-specific genomic response in D. melanogaster populations during experimental evolution in hot and cold environments. Mol Biol Evol 31: 364–375.

True, J. R., J. M. Mercer and C. C. Laurie, 1996 Differences in crossover frequency and distribution among three sibling species of Drosophila. Genetics 142: 507–523.

Turner, T. L., and P. M. Miller, 2012 Investigating natural variation in Drosophila courtship song by the evolve and resequence approach. Genetics 191: 633–642.

Turner, T. L., A. D. Stewart, A. T. Fields, W. R. Rice and A. M. Tarone, 2011 Population-based resequencing of experimentally evolved populations reveals the genetic basis of body size variation in Drosophila melanogaster. Plos Genetics 7.

Vlachos, C., and R. Kofler, 2018 MimicrEE2: Genome-wide forward simulations of Evolve and Resequencing studies. PLOS Computational Biology 14: e1006413.

Wallace, B., 1970 Genetic Load. Its Biological and Conceptual Aspects. Prentice-Hall, Englewood Cliffs, N.J.

Walsh, B., and M. Lynch, 2018 Evolution and Selection of Quantitative Traits. http://nitro.biosci.arizona.edu/zbook/NewVolume2/pdf/Chapter09.pdf.

Ward, J. K., and J. K. Kelly, 2004 Scaling up evolutionary responses to elevated CO_2_: lessons from Arapidopsis. Ecology Letters 7: 427–440.

Wright, S., 1931 Evolution in mendelian populations. Genetics 16: 97–159.

Wright, S., 1943 Isolation by distance. Genetics 28: 114.

